# Combinatorial proteomics and transcriptomics identify AMPK in the control of the axonal regeneration programme of DRG sensory neurons after spinal injury

**DOI:** 10.1101/661488

**Authors:** Guiping Kong, Luming Zhou, Elisabeth Serger, Ilaria Palmisano, Francesco De Virgiliis, Thomas H Hutson, Eilidh Mclachlan, Anja Freiwald, Paolo La Montanara, Kirill Shkura, Radhika Puttagunta, Simone Di Giovanni

## Abstract

Regeneration after injury occurs in axons that lie in the peripheral nervous system but it fails in the central nervous system limiting functional recovery. Despite recent progress, the signalling response to injury of peripheral versus central projecting axons that might underpin this differential regenerative ability is currently largely uncharacterized. To fill this knowledge gap, here we combined axoplasmic proteomics from peripheral sciatic or central projecting dorsal root axons from sciatic DRG neurons. Proteomics was combined with cell body RNAseq to compare axonal and soma responses between a spinal regeneration-incompetent versus sciatic regeneration-competent nerve injury. This allowed the identification of injury-dependent signalling pathways uniquely represented in peripheral versus central projecting sciatic DRG axons. RNAseq and proteomics analysis suggested AMPK as a putative regulator of axonal regenerative signalling pathways. AMPK immunoprecipitation followed by mass spectrometry from DRG suggested that the 26S proteasome and its regulatory subunit PSMC5 preferentially interact with AMPKα for proteasomal degradation following sciatic axotomy. Mechanistically, we found that the proteasome and CaMKIIα-dependent proteasomal subunit PSMC5 regulates AMPKα1 protein expression. Finally, conditional deletion of AMPKα1 promoted multiple regenerative signalling pathways and robust axonal growth across the injured spinal cord, suggesting inhibition of AMPK as a novel regenerative target following spinal injury.

**HIGHLIGHTS:**

- Axoplasmic proteomics from sciatic or centrally projecting branches of sciatic DRG identifies unique protein enrichment and signalling pathways, including prior and subsequent to a spinal regeneration-incompetent versus sciatic regeneration-competent axonal injury
- Combined axoplasmic DRG proteomics and cell body RNAseq analysis suggest AMPK as a central regulator controlling axonal regeneration
- The 26S proteasome and the 19S regulatory subunit PSMC5 interact with AMPKα following sciatic axotomy. PSMC5 regulates AMPKα1 protein levels
- AMPKα1 conditional deletion enhances robust axonal growth following SCI

## INTRODUCTION

While axonal regeneration and partial functional recovery in the injured peripheral nervous system (PNS) occur, these fail in the central nervous system (CNS) such as after a spinal cord injury (SCI) ^1^. The pseudounipolar sensory fiber tracts belonging to the dorsal root ganglia (DRG) system extending one branch in the periphery and one into the central spinal column represent a fruitful model for direct comparison of the differential regenerative capacities of PNS and CNS axons sharing a single cell body. Although leading to limited axonal regeneration beyond the lesion site, the gold standard for regeneration of sensory fibers across the injured spinal cord remains the conditioning lesion of the peripheral sciatic nerve preceding a spinal cord injury ^2,3^.

Assessment of the molecular mechanisms that drive sciatic nerve regeneration has revealed that in response to injury, signals are transmitted from the environment, transduced via cell surface receptors, such as IL-6R, tyrosine kinase receptors, and transported through retrograde signalling to the soma of the DRG ^4^. Within the DRG neuron, these events promote the activation of intracellular signalling cascades and proteins, such as the dual leucine zipper kinase (DLK)^5^, mitogen-activated protein kinase pathway (MAPK) and phosphoinositide 3-kinase (PI3 K) that culminate in the activation of regeneration-specific transcription factors (TFs) such as pSTAT3, c-JUN, pCREB, ATF3^6^. The activity of these and other transcription factors are in turn modulated by epigenetic co-factors including p300, CBP, PCAF and HDAC5^7–11^, leading to the simultaneous induction of a large number of regenerative associated genes (RAGs). Attempts to stimulate the intrinsic regenerative response within the injured CNS have been only partially successful through the overexpression/knockdown of individual genes and TFs, including c-JUN, pCREB, SMAD1, PTEN, KLF4, KLF7 and STAT3^12–18^.

Pathways related to protein synthesis and energy metabolism such as PTEN-mTOR signalling are also essential to regulate axonal regeneration of the corticospinal tracts, the optic nerve as well as central sensory DRG axons. In fact, deletion of PTEN and activation of mTOR strongly contribute to regeneration of these fiber tracts ^15,19,20^. Therefore, it is evident that both transcriptional control and protein synthesis are required for the axonal regeneration programme. These studies also suggest that axonal trafficking following peripheral but not central axonal injury regulates pathways that positively fine-tune the regenerative phenotype, including the modulation of long-term nuclear changes responsible for regenerative reprogramming.

However, despite this recent progress, we are still uncovering the nature of the contrasting molecular signatures associated with successful PNS versus failed CNS axonal regeneration, limiting the identification of effective regenerative targets. Additionally, the centrally and peripherally projecting axons originating from sciatic DRG seem to display a differential axonal trafficking, possibly reflecting a distinct energy metabolism, microtubule and actin cytoskeleton^21^.

However, systematic studies comparing how these two compartments respond to injury are still missing. Here, we performed proteomics from the axoplasm of sciatic nerve or centrally projecting axons (dorsal roots) to investigate the molecular signatures associated with a regenerative-competent versus a regenerative-incompetent axonal compartment both prior and subsequent to an equidistant sciatic or spinal cord (dorsal column) axotomy. Axoplasmic proteomics following sciatic or spinal injury showed that the expression of the α catalytic subunit of AMPK was significantly reduced by a sciatic injury while it remained unchanged in the central projecting DRG branches following SCI.

AMPK is a central regulator of metabolic signalling and its activity is typically controlled by ATP, ADP and AMP levels ^22–25^. AMPK phosphorylates a number of enzymes directly involved in these processes as well as transcription factors, co-activators, and co-repressors affecting transcription and intracellular signalling, including insulin, JAK-STAT and mTOR dependent pathways, previously implicated in axonal regeneration ^26–28^. We next integrated the proteomics profiles with recently generated RNAseq from sciatic DRG cell bodies ^29^. This combined analysis identified several metabolic and signalling pathways involved in ATP metabolism, glucose and protein synthesis as well as in insulin and mTOR pathways among others. This further suggested a potential novel role for AMPK in controlling the axonal regeneration programme. Therefore, next we confirmed that AMPKα1 protein expression as well as its active phosphorylated form were reduced following sciatic nerve injury. In order to identify possible AMPK interacting proteins responsible for AMPKα downregulation, we performed AMPKα immunoprecipitation followed by mass spectrometry from DRG *ex vivo* following sciatic injury. Bioinformatics analysis suggested the 26S regulatory proteasomal subunit PSMC5 as novel AMPKα1 interacting partner. Indeed, we found by co-immunoprecipitation that AMPKα1 and PSMC5 interact and that post-injury AMPKα1 levels depends upon CaMKIIα, which is known to phosphorylate and activate PSMC5 ^30^. Importantly, we discovered that AMPKα1 controls the expression of the regenerative programme since conditional deletion of AMPKα1 in sciatic DRG neurons *in vivo* activates several members of signalling pathways associated with axonal regeneration. Finally, conditional deletion of AMPKα1 but not α2 promoted significant axonal regeneration of sensory ascending DRG axons across the injured spinal cord, suggesting inhibition of AMPK as a novel regenerative target following spinal injury.

## RESULTS

### Combinatorial axoplasmic proteomics and RNAseq from DRG suggest AMPK as a putative regulator of differential regenerative ability between central and peripheral injury

Our initial experiments aimed to systematically investigate the expression of previously unknown axonal proteins and signalling pathways associated with regeneration-competent sciatic nerve vs regeneration-incompetent central projecting axonal branches. This included analysis in sham injury control conditions in comparison to peripheral sciatic nerve or spinal dorsal column axotomy. We performed protein mass spectrometry from axoplasmic extracts from three independent biological replicates of sciatic nerve or central projecting axonal branches (Fig. 1a, b). We used stable isotopes quantitative proteomics methodology, which allows evaluation of peptide relative abundances between two samples (sample and control) by labelling them with reagents that contain either a heavy and or a light mass isotope. After mixing the two, the labelled samples are analysed by mass spectrometry. The peaks in the mass spectra reveal the ratio of the two different isotopes that is a measure of the peptide relative abundances^31^. This allowed the identification of a total of 3128 protein groups. In recent previous experiments, we have carried out RNAseq from the soma of sciatic DRG following the same dual injury paradigm including a DRG equidistant sciatic nerve axotomy and a spinal dorsal column axotomy ^29^. Here, we combined this previous gene expression analysis with our newly acquired axoplasm proteomics data (Fig. 1a, b).

**Figure 1.**
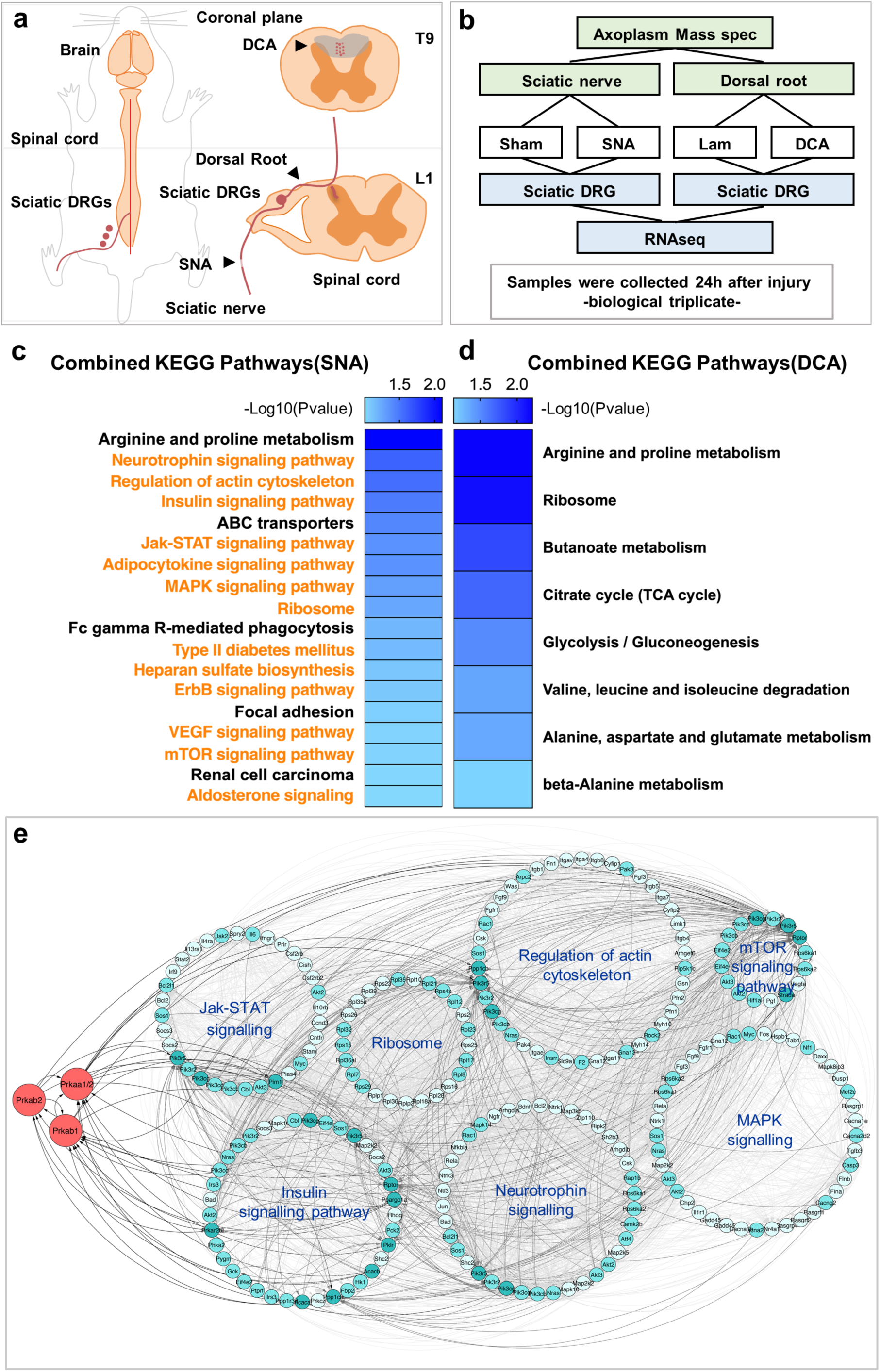
Combined proteomics and RNAseq data analysis identify AMPK as a potential regulator of axonal regeneration. **a**, Diagram of sciatic nerve and T9 dorsal column axotomy. **b**, Schematic diagram of the experimental paradigm. Axoplasm Mass Spectrometry was performed in biological and technical triplicate. **c**, **d**, Heatmap graphs show the enriched KEGG pathways of the differentially expressed genes (p-value < 0.05) and proteins (FDR < 0.05, |log2ratio|> 0.58), in DRG and axoplasm respectively, after SNA (c) or DCA (d). KEGG pathways in orange are pathways that have been found to be regulated by AMPK. **e**, Cytoscape visualization of the protein network under AMPK control after SNA with combined analysis of proteomics and RNAseq data. Protein-protein interaction network was generated by String and visualized by Cytoscape. The nodes represent individual proteins, while edges represent protein interaction score according to String database. Proteins were jointly organized in a circular layout according to their KEGG affiliation by using Cytoscape. Red nodes represent the three AMPK subunits.

First, we characterized the degree of neuronal enrichment versus other neighbouring cell types of the axoplasmic proteins identified by mass spectrometry. As expected, axoplasmic proteins with neuronal identity were highly enriched compared to protein expressed selectively by Schwann cells or oligodendrocytes. This was measured by assessing how many of our detected protein groups were shared by single cell DRG neuronal versus Schwann cell or oligodendrocyte specific RNAseq or microarray gene expression studies ^32–34^. This analysis allowed us to establish that only 277 out of 2348 proteins identified in our peripheral axoplasmic preparation and 204 out of 2143 in central axoplasm, representing about 12% and 10% of the total, were matched for gene expression in Schwann cells but not in DRG neurons (Supplementary Fig. 1a). Only 63 and 56 proteins in both peripheral and central axoplasm, representing about 3% of the total, were also found in oligodendrocytes but not in DRG neurons (Supplementary Fig. 1b).

In search for signalling pathways and master regulators able to discriminate between axonal regeneration and regenerative failure, we compared proteomics expression data of injured peripheral and central axons (following sciatic nerve axotomy-SNA or spinal dorsal column axotomy –DCA respectively) vs their corresponding sham controls. We found 172 and 140 differentially expressed proteins following SNA or DCA respectively (1.5-fold up- or down-regulated, FDR < 0.05, Supplementary File 1). Interestingly, unsupervised clustering (Supplementary Fig. 2a) and direct comparison of differentially expressed proteins following SNA or DCA (Supplementary Fig. 2b) showed that the two injury paradigms were associated with two very distinct protein expression profiles (Supplementary File 1). Gene ontology (GO) analysis of the proteins specifically downregulated following SNA showed enrichment for ribosomal protein synthesis and translation, structural molecule activity, actomyosin cytoskeleton, ATP-dependent metabolism and regulation as well as in adenyl nucleotide binding (Supplementary Fig. 2c, Supplementary File 3). Remarkably, all of these functional classes (Supplementary Fig. 2c, highlighted in orange) are well known to regulate and be regulated by AMPK ^22,35–39^. Additionally, these signalling pathways were recently found to be enriched in the proteome interacting with AMPKα following immunoprecipitation of AMPKα and mass spectrometry analysis in non-neuronal cells^40^. Importantly, the analysis of the proteins belonging to these functional classes revealed that AMPKα, the catalytic subunit of the AMPK complex, is significantly downregulated following SNA compared to sham (log ratio: −0.89 and FDR < 0.05, Supplementary File 1). Proteins groups specifically overexpressed in the sciatic nerve after nerve injury were mainly enriched for cell adhesion, transcription regulation, and carbohydrate metabolism (Supplementary Fig. 2d, Supplementary File 2). A number of previously characterized regeneration-associated genes (RAGs) were also found enriched at the protein level in the axoplasm following sciatic but not spinal injury, supporting the robustness of our data (Supplementary Fig. 3a and Supplementary File 1). Proteins differentially expressed following DCA were mainly associated with mitochondrial function (upregulated) and protein synthesis (downregulated) (Supplementary Fig. 2e, f). We next performed validation of the axoplasmic proteomic dataset by immunoblotting of purified axoplasm following SNA or DCA versus sham or laminectomy controls respectively. This confirmed expression changes of several proteins, including RAGs (Supplementary Fig. 3a) and allowed assessing the degree of enrichment and purity of our axoplasmic preparation by comparing it to protein extraction from the spinal cord, sciatic nerve and DRG (Supplementary Fig. 3b-e). We found that the peripheral axoplasm showed only low expression of S100 (Schwann cell marker) and myelin protein zero (MPZ) compared to the sciatic nerve and DRG (Supplementary Fig. 3b, c). The central axoplasm was devoid of detectable levels of MOG (myelin oligodendrocyte glycoprotein) and S100 as compared to high expression in the spinal cord and DRG (S100) (Supplementary Fig. 3d, e).

In order to gain a better understanding of the signalling pathways within the “DRG axonal signalling unit” that may be relevant for the regenerative transcriptional programme, we combined the axoplasmic proteomics and previously published DRG transcriptomics data ^29^ by creating a reference data set, which can be used for a joint pathway analysis following standard methodologies and as further described in the methods ^41,42^. We performed KEGG pathway analysis of the combined RNAseq and proteomics datasets that showed enrichment in previously described regenerative signalling as well as in metabolic signalling pathways, ribosomal protein synthesis, and in actin regulation following SNA (Fig. 1c and Supplementary File 3). Interestingly, several of these signalling pathways previously implicated in axonal regeneration have been found to be regulated by AMPK. They include neurotrophin, insulin, mTOR, JAK-STAT, MAPK, ErbB and VEGF signalling ^26–28,43–46^, while others are related to AMPK-dependent metabolic regulation such as carbohydrate, amino acid metabolism ^26,47–50^ or adipocytokine signalling ^51,52^.

However, following DCA, fewer genes and proteins were differentially regulated and they were mainly represented by amino acid and catabolic metabolism (Fig. 1d and Supplementary File 3).

Together, the data so far suggest that AMPK might control multiple signalling pathways that are involved in axonal regeneration. This is further emphasized by a network analysis representing the functional connections between AMPK and signalling pathways associated to differential expressed genes and proteins implicated in regenerative signalling (Fig. 1e).

### AMPKα1 expression, phosphorylation and activity are downregulated following SNA

Axoplasmic proteomics indicated that AMPKα protein expression was significantly reduced following SNA versus sham, but remained unchanged following DCA versus laminectomy (Supplementary File 1). Before investigating AMPKα expression and activation further, we explored whether reduction in AMPKα expression in the axoplasm after SNA was or not restricted to one of the two AMPK catalytic subunits α1 and α2. Immunoblotting confirmed a reduction in AMPKα1 but not α2 expression in the axoplasm following SNA (Fig. 2a, b). Next, we aimed to establish the expression levels of AMPKα1 and α2 following central DCA versus peripheral SNA in sciatic DRG. Immunoblotting revealed that the expression of AMPKα1 but not α2 is reduced following SNA and it remained unchanged after DCA (Fig. 2c-f). While the expression of the overall active phosphorylated AMPKα (p-AMPKα) is also reduced following SNA, p-AMPKα is diminished to a similar extent as total AMPKα1 (Fig. 2c-f), suggesting that sciatic axotomy is followed by reduction of AMPKα1 protein expression rather than specific changes in phosphorylation. In order to detect whether reduced AMPK levels also affects AMPK activity, we measured by ELISA AMPKα activity from DRG homogenates following SNA or DCA after addition of ATP by using insulin receptor substrate-1 S789 (IRS)-1 S789 as substrate (see methods for details). We found that AMPK activity is significantly reduced after SNA but not DCA compared to sham control (Fig. 2 g, h), in line with the reduction of AMPKα1 and p-AMPKα protein expression. Importantly, expanding upon the immunoblotting and activity data, we also found that AMPKα1 but not AMPKα2 expression was significantly decreased in NF200^+^ DRG neurons following SNA (Fig. 2i-n), however neither AMPKα1 nor AMPKα2 expression was affected by DCA (Fig. 2i-n). Interestingly, while AMPKα1 signal showed a cytoplasmic localisation, AMPKα2 was mainly nuclear, suggesting independent functions in DRG neurons.

**Figure 2.**
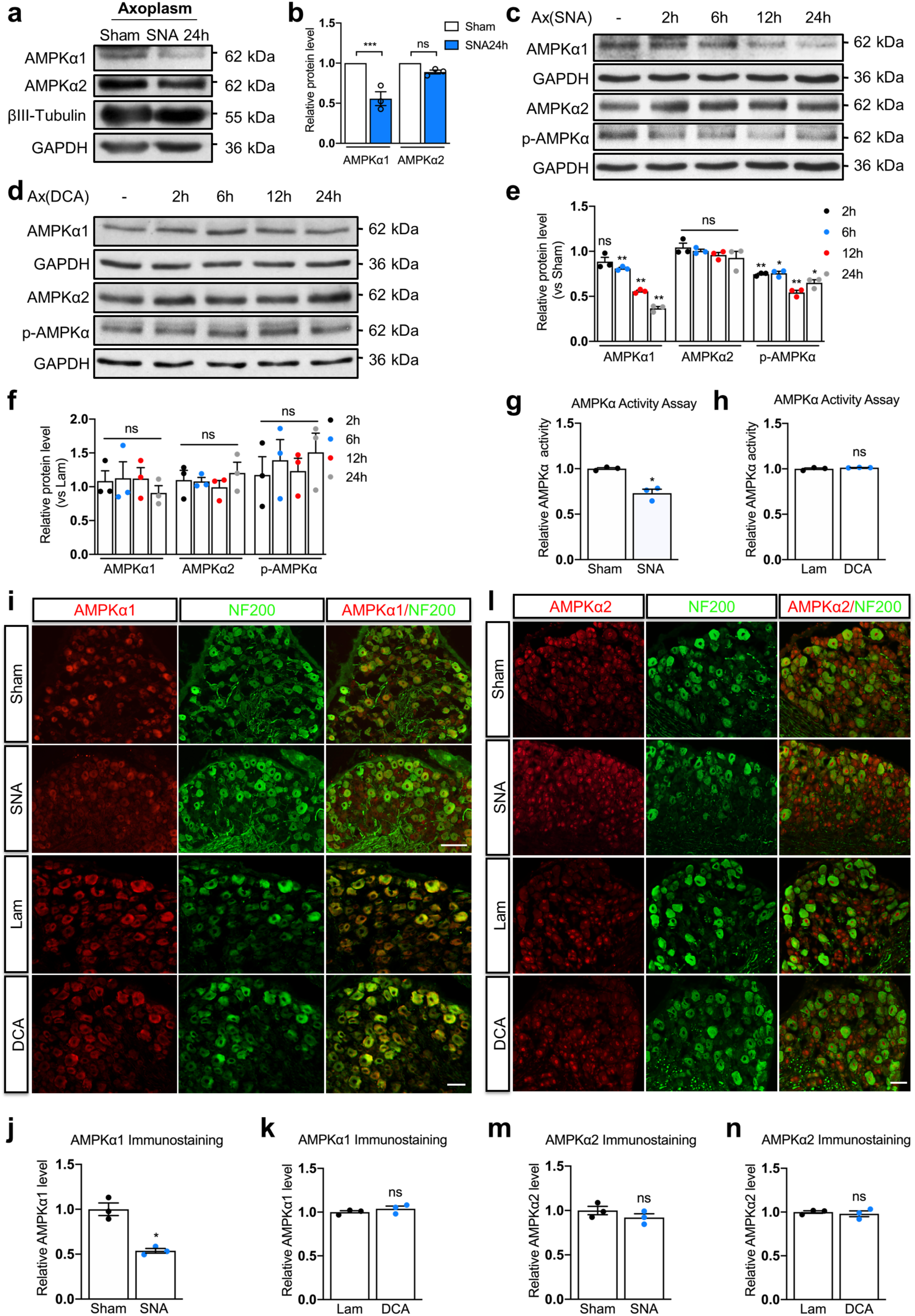
AMPKα1 expression and activity are downregulated following SNA. **a**, Immunoblot shows a reduction of AMPKα1 but not of AMPKα2 expression in the sciatic axoplasm following SNA at 24 h. **b**, Immunoblot quantification of AMPKα1 and AMPKα2 expression levels in (**a**) (n = 3 independent experiments). The relative protein expression level was quantified after normalization to GAPDH. Values represent means ± SEM (***p < 0.001; ns: not significant; two-way ANOVA followed by Bonferroni test). **c**, **d**, Immunoblots show AMPKα1; AMPKα2 and p-AMPKα expression in sciatic DRGs after SNA (**c**) or DCA (**d**) at several time points. **e**, **f**, Immunoblots quantification of AMPKα1; AMPKα2 and p-AMPKα expression levels show reduction of AMPKα1 and p-AMPKα following SNA but not DCA (n = 3 independent experiments). The relative expression level of each protein at individual time point was quantified after normalization to GAPDH. Values represent means ± SEM (*p < 0.05; **p < 0.01; ns: not significant; two-way ANOVA followed by Bonferroni test). **g**, **h**, AMPK activity assay from DRG (n = 3 independent experiments). The assay was performed from sciatic DRG samples following Sham or SNA (24 h) (**g**) and Lam or DCA (24 h) (**h**). Values represent means ± SEM (*p < 0.05; ns: not significant; two-tailed unpaired Student’s *t*-test). **i**, **l**, Representative fluorescence images of immunostaining for AMPKα1 (**i**) or AMPKα2 (**l**); neurofilament 200 (NF-200) in DRG neurons following Sham or SNA (24 h); Lam or DCA (24 h). In (**i**): upper scale bar, 100 μm; lower scale bar, 50 μm. In (**l**): scale bar, 50 μm. **j**, **k**, **m**, **n**, Quantification of immunofluorescence intensity for AMPKα1 (**j**, **k**) or AMPKα2 (**m**, **n**) (n = 3). The relative AMPKα1 and α2 expression levels were quantified after normalization to the background immunofluorescence (secondary antibody only). Values represent means ± SEM (*p < 0.05; ns: not significant; two-tailed unpaired Student’s *t*-test).

We next investigated the expression of AMPKα1 in DRG neuron subtypes by co-immunohistochemistry experiments. We found that AMPKα1 is strongly expressed in the large majority of NF200 and Parvalbumin (PARV) positive DRG mechanoceptive and proprioceptive neurons, while it is poorly expressed in peptidergic (CGRP positive) and non-peptidergic (IB4 positive) nociceptors (Supplementary Fig. 4a, b). Expression of AMPKα1 also decreased in PARV^+^ DRG neurons following SNA (Supplementary Fig. 4c, d), while no significant expression changes were observed in nociceptors (n = 3, non-significant change in signal intensity in sham vs SNA, two-tailed *t*-test).

### AMPKα1 controls DRG regenerative growth

To investigate whether inhibition of AMPK activity would promote DRG neurite outgrowth, we used the AMPK inhibitor compound C (an ATP competitor) in cultured DRG neurons. We found that pharmacological inhibition of AMPK activity with compound C (10nM) significantly enhanced neurite outgrowth in cultured DRG neurons on both PDL/laminin growth permissive and myelin growth inhibitory substrates (Fig. 3a, b). Since it has been reported that AMPK regulates cell growth via suppression of the mammalian target of rapamycin complex 1 (mTORC1) pathway ^53^, we tested whether compound C would enhance the phosphorylation of ribosomal protein S6 kinase (p70S6 K), which is a downstream target of mTOR ^54^. Immunoblotting showed that treatment with 10nM compound C increased p70S6 K expression (Fig. 3c, d), supporting the evidence for inhibition of AMPK signalling. Next, we measured neurite outgrowth after compound C or vehicle following conditional deletion of AMPKα1 by AAV-cre-GFP or AAV-GFP delivery in cultured DRG neurons from AMPKα1 floxed mice. This experiment showed that conditional deletion of AMPKα1 promotes neurite outgrowth and that compound C does not alter the phenotype associated with AMPKα1 deletion (Fig. 3e, f), suggesting that compound C mediated increase in outgrowth relies upon AMPKα1 deletion. Immunofluorescence for AMPKα1 confirmed strong reduction of AMPKα1 expression (Supplementary Fig. 5a and Fig. 3 g). Lastly, as direct proof that AMPKα1 restricts neurite outgrowth of DRG neurons, we overexpressed an AMPKα1 or GFP plasmid DNA in DRG neurons to find that AMPKα1 overexpression represses neurite outgrowth (Fig. 3h-j). We also did not observe any difference in cell density between AMPKα1 and GFP plasmid transfected cells as well as in apoptotic features as confirmed by the analysis of the nuclear morphology by DAPI staining (Supplementary Fig. 5b, c).

**Figure 3.**
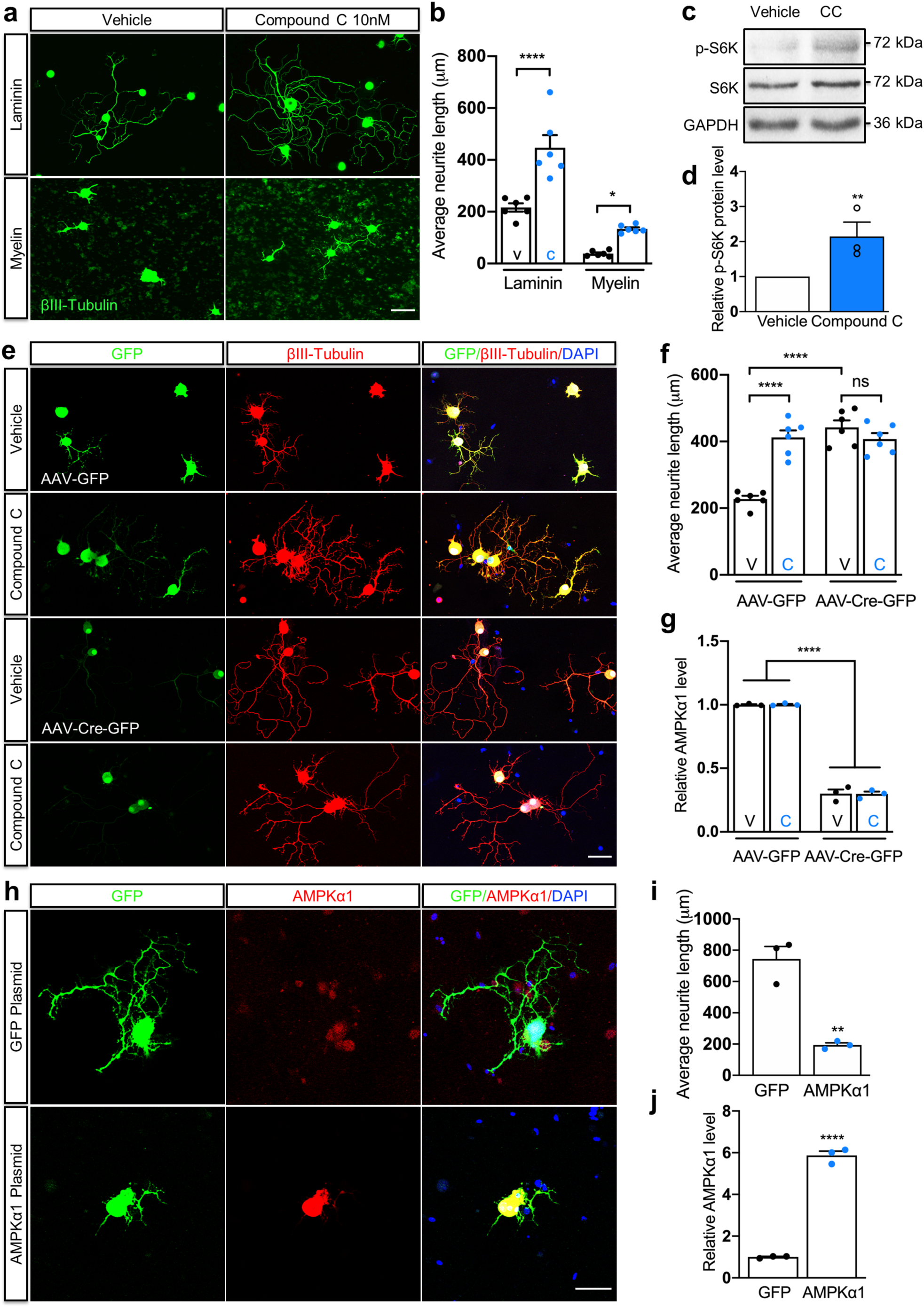
AMPKα1 inhibits DRG regenerative growth. **a**, Representative neurite outgrowth images of cultured DRG neurons 24 h after delivery of vehicle or compound C (AMPK activity inhibitor). Compound C promotes DRG regenerative growth. Scale bar, 50 μm. **b**, Quantification of average neurite length of (**a**). V represents vehicle, C represents compound C. n = 3 independent experiments in technical replicate. Values represent means ± SEM (*p < 0.05; ****p < 0.0001; two-way ANOVA followed by Bonferroni test). **c**, Immunoblot shows p-S6 K and S6 K expression in cultured DRG neurons treated with vehicle or compound C (10nM). **d**, Immunoblot quantification of p-S6 K expression level of (**c**). n = 3 independent experiments. The relative protein level was quantified following normalization to GAPDH. Values represent means ± SEM (**p < 0.01; two-tailed unpaired Student’s *t*-test). **e**, Representative neurite outgrowth images of cultured DRG neurons 48 h after transfection with AAV-GFP or AAV-Cre-GFP. DRG neurons were dissected from AMPKα1 floxed mice and dissociated DRG cells were treated with vehicle or 10nM compound C together with GFP or Cre virus. Scale bar, 50 μm. **f**, Quantification of average neurite length of (**e**). V represents vehicle, C represents compound C. n = 3 independent experiments in technical replicate. Values represent means ± SEM (****p < 0.0001; ns: not significant; two-way ANOVA followed by Bonferroni test). **g,** Quantification of AMPKα1 expression level of (**e**). Representative fluorescence images of AMPKα1 staining of (**e**) are provided in Supplementary Fig. 5a. V represents vehicle, C represents compound C. n = 3 independent experiments. Values represent means ± SEM (****p < 0.0001; two-way ANOVA followed by Bonferroni test). **h**, Representative neurite outgrowth images of cultured DRG neurons 48 h after electroporation with GFP or AMPKα1 plasmid. Scale bar, 100 μm. **i**, Quantification of average neurite length of (**h**). n = 3 independent experiments. Values represent means ± SEM (**p < 0.01; two-tailed unpaired Student’s *t*-test). **j**, Quantification of AMPKα1 expression level of (**h**). n = 3 independent experiments. Values represent means ± SEM (****p < 0.0001; two-tailed unpaired Student’s *t*-test).

Together, these data indicate that the expression of AMPKα1 and AMPK activity are reduced following regeneration-competent SNA while they remain unchanged compared to control following regeneration-incompetent DCA. Inhibition of AMPK activity promotes DRG neurite outgrowth *in vitro* on both growth permissive and inhibitory substrates and relies on AMPKα1. Lastly, AMPKα1 conditional deletion or overexpression enhance or reduce outgrowth of DRG neurons respectively, implicating AMPKα1 in the control of regenerative growth of sensory neurons.

### AMPKα immunoprecipitation and mass spectrometry indicate that AMPKα associates with the proteasome, which regulates AMPKα protein levels after peripheral nerve injury

To investigate the molecular mechanisms underpinning the SNA-dependent reduction of AMPKα1 protein expression, we performed immunoprecipitation of AMPKα followed by mass spectrometry from DRG ex vivo following sham or sciatic nerve axotomy. We used stable isotopes quantitative proteomics methodology, where AMPK co-interactors were identified in either sham or SNA condition vs IgG that was used as a negative control to exclude the non-specifically bound proteins. While as expected AMPKα was one of the most represented proteins, we found that 172 and 94 proteins immunoprecipitated with AMPKα following sham or sciatic injury respectively (FDR < 0.05, mean log2 ratio > 0.58, Supplementary File 4). However, statistical analysis of differentially enriched proteins showed that 33 proteins immunoprecipitated uniquely or preferentially (Fisher FDR < 0.05, FC SNA vs Sham, >1.5, Supplementary File 4) upon SNA, and 78 immunoprecipitated uniquely or preferentially after sham (Fisher FDR < 0.05, FC SNA vs Sham, < 0.5, Supplementary File 4).

To gain further functional insight, we performed a protein-protein interaction network analysis (using String followed by Cytoscape) of proteins identified by AMPKα IP-mass spec. Network analysis of the AMPKα interactome showed that multiple proteins interacting with AMPKα belonged to proteasomal and ribosomal protein networks (Fig. 4a). This might explain the fact that downregulated proteins in the axoplasm were enriched for functional categories related to protein translation (Supplementary File 2). Interestingly, we found an increased affinity of AMPKα with several subunits of the 26S proteasome following SNA (Fig. 4a and Supplementary File 4).

**Figure 4.**
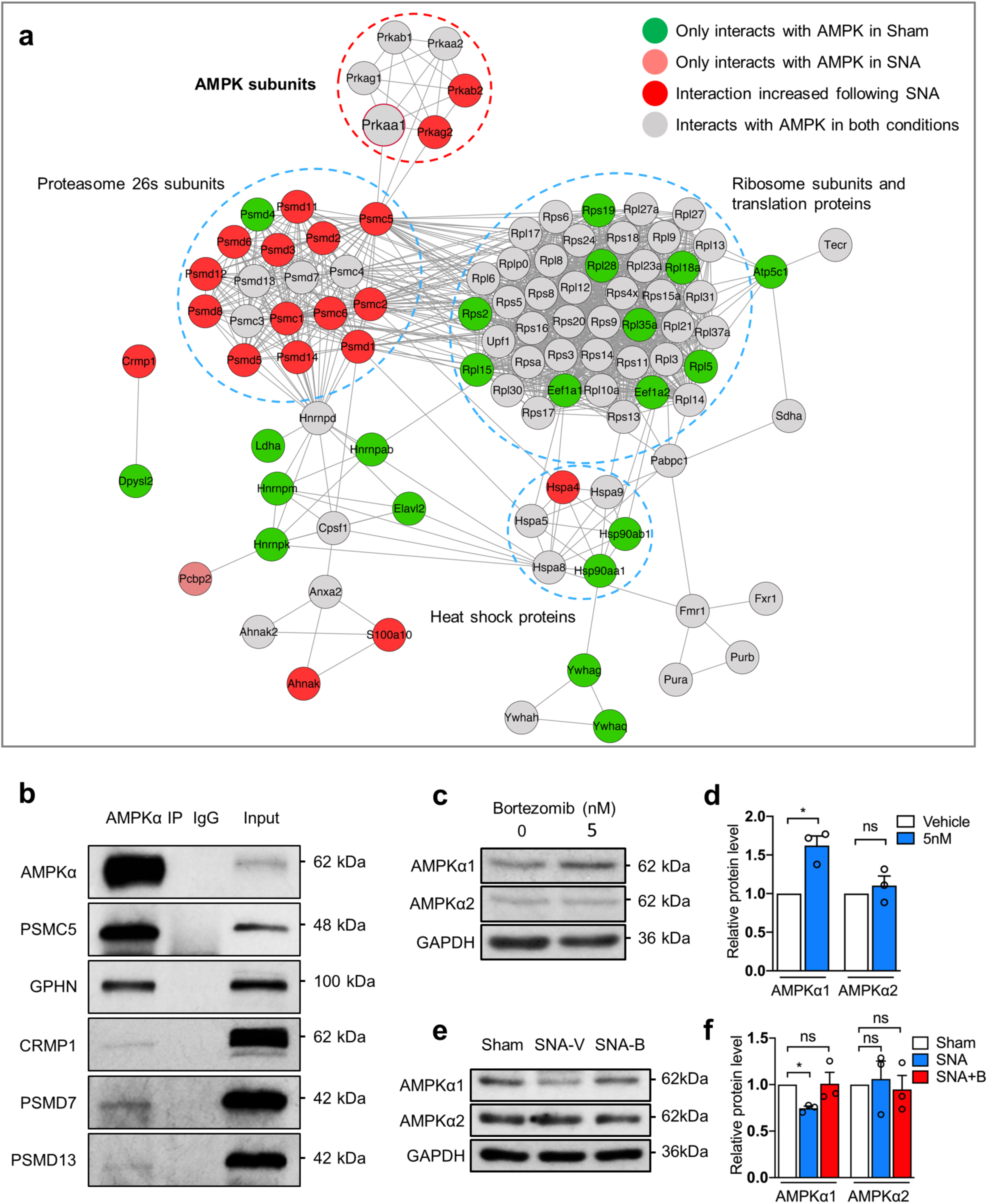
AMPKα interacts with the proteasome including the proteasomal subunit PSMC5 and AMPKα1 expression is regulated by proteasome activity. **a**, Network analysis of AMPKα protein interactions identified by AMPKα IP-mass spec following SNA vs Sham. Proteins were selected based upon FDR < 0.05, mean log2 ratio >0.58 (Sham vs IgG or SNA vs IgG). n = 2 replicates. Protein network was generated by String and visualized by Cytoscape. The nodes represent individual proteins, while edges represent interaction scores. Node colour defines the interaction with AMPK following SNA and Sham (light red: only interacts with AMPK after SNA; dark red: increased interaction after SNA, Fisher FDR < 0.05, FC (SNA vs Sham) >1.5; green: only interacts with AMPK in Sham condition; grey: interacts with AMPK in both Sham and SNA). **b**, Immunoblot shows the validation of candidates following AMPKα IP in DRG neurons. These candidates were chosen based on their log2 ratios and FDR in AMPK IP MS dataset. **c**, Immunoblot shows AMPKα1 and AMPKα2 expression in F11 cells after incubation with 5nM Bortezomib (proteasome inhibitor) at 6 h. **d**, Quantification of AMPKα1 and AMPKα2 expression levels of (**c**). n = 3 independent experiments. Relative protein expression level was quantified after normalization to GAPDH. Values represent means ± SEM (*p < 0.05; ns: not significant; two-way ANOVA followed by Bonferroni test). **e**, Immunoblot shows AMPKα1 and AMPKα2 expression in DRG after intraperitoneal injection of Bortezomib (1 mg/kg, 3 times i. p. injections per day) 24 h following SNA. **f**, Quantification of AMPKα1 and AMPKα2 expression levels of (**e**). n = 3 independent experiments. Relative protein expression level was quantified after normalization to GAPDH. Values represent means ± SEM (*p < 0.05; ns: not significant; two-way ANOVA followed by Bonferroni test).

Among these subunits, the 19S regulatory subunit PSMC5, also known as RPT6, is predicted to connect AMPK with the proteasome (Fig. 4a), although experimental evidence suggesting the formation of a direct protein interaction is not available to date. Together, this allowed us to formulate the hypothesis that SNA-dependent reduction in AMPKα1 expression might be mediated by increased proteasome degradation through the 26S proteasome and that the 19S regulatory subunit PSMC5 might be a critical component. Indeed, PSMC5/RPT6 has been reported as a crucial subunit for the proteasome 19S regulatory particle assembly and proteasome activation ^55^.

Therefore, we asked whether PSMC5 interacts with AMPKα by performing co-immunoprecipitation experiments and immunoblotting in DRG. In line with our hypothesis, we found that AMPKα co-immunoprecipitated with PSMC5 (Fig. 4b). We further validated the identification AMPKα interacting proteins by IP-mass spectrometry by performing IP for AMPKα and immunoblotting for the axonal growth protein CRMP1, postsynaptic protein GPHN and for two additional proteasomal proteins PSMD7 and PSMD13 belonging to the PSMC5 protein cluster. These proteins indeed co-immunoprecipitated with AMPKα although with an apparent lower affinity than PSMC5 (Fig. 4b). By performing immunoblotting for PSMC5 from DRG following immunoprecipitation using antibodies specific to AMPKα1 or α2, we found that both subunits co-immunoprecipitated with PSMC5 (Supplementary Fig. 6a, b).

Next, we investigated whether the 26S proteasome controls AMPKα1 and α2 protein level. We performed immunoblotting of AMPKα1 and α2 from F11 DRG cell lines and from DRG *ex vivo* after *in vitro* or *in vivo* i.p. delivery of the 26S proteasome inhibitor Bortezomib respectively. Bortezomib administration enhanced AMPKα1 but not α2 protein expression both in culture and *ex vivo,* including after sciatic nerve axotomy (Fig. 4c-f), suggesting that the 26S proteasome affects AMPKα1 protein levels.

### PSMC5 regulates AMPKα1 expression and neurite outgrowth of DRG neurons

In order to confirm that PSMC5 regulates AMPKα1 protein expression, which is affected by manipulation of the proteasome, we performed gene silencing of PSMC5 after transfection with PSMC5 siRNA in F11 DRG cell lines and measured AMPKα1 protein expression via immunoblotting to find that AMPKα1 was significantly up-regulated following PSMC5 silencing (Fig. 5a, b). Next, we electroporated primary cultured DRG neurons with the same siRNA-GFP against PSMC5 vs a scrambled control siRNA-GFP used in F11 DRG cells - to verify whether it would inhibit neurite outgrowth. Indeed, we found that PSMC5 gene silencing inhibited neurite outgrowth in individual GFP positive neurons showing reduced PSMC5 expression (Fig. 5c-e). We next investigated whether inhibition of AMPK activity would overcome reduction of neurite outgrowth following PSMC5 silencing. We delivered compound C as described in earlier experiments (Fig. 3) following siRNA-GFP against PSMC5 vs a scrambled control siRNA-GFP in primary cultured DRG cells. Data analysis revealed that compound C promoted neurite outgrowth even in the presence of enhanced AMPKα1 expression following PSMC5 silencing (Fig. 5f-h).

**Figure 5.**
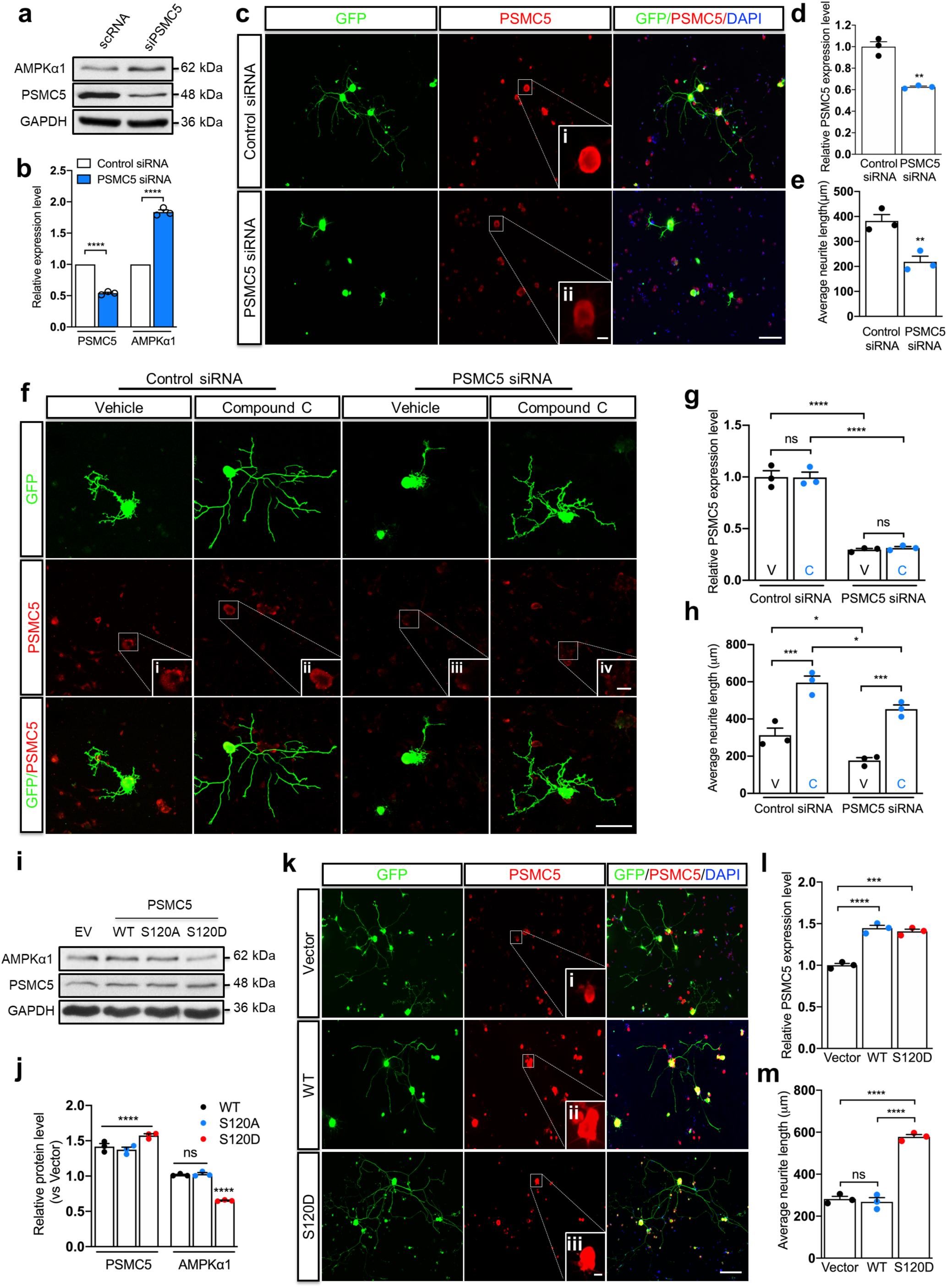
PSMC5 is required for AMPKα1 expression levels. **a**, Immunoblot shows PSMC5 and AMPKα1 expression 48 h after transfection with control siRNA or PSMC5 siRNA in F11-DRG cell lines. **b**, Quantification of PSMC5 and AMPKα1 expression levels of (**a**). n = 3 independent experiments. Relative protein expression level was quantified after normalization to GAPDH. Values represent means ± SEM (****p < 0.0001; two-way ANOVA followed by Bonferroni test). **c**, Representative neurite outgrowth images of cultured DRG neurons 36 h after electroporation with control siRNA or PSMC5 siRNA. Cells were immunostained with anti-PSMC5, anti-GFP and counterstained with DAPI. Scale bar, 200 μm. **c** (i-ii), scale bar, 20 μm. **d**, Quantification of PSMC5 expression level of (**c**). n = 3 independent experiments. Relative PSMC5 expression level in siRNA transfected DRG cells was quantified versus control scRNA. Values represent means ± SEM (**p < 0.01; two-tailed unpaired Student’s *t*-test). **e,** Quantification of average neurite length of (**c**). n = 3 independent experiments. Values represent means ± SEM (**p < 0.01; two-tailed unpaired Student’s *t*-test). **f**, Representative neurite outgrowth images of cultured DRG neurons 36 h after electroporation with control siRNA or PSMC5 siRNA. Vehicle and 10nM compound C were added in fresh cell culture medium 12 h after electroporation. Scale bar, 100 μm. **f**(i-iv), scale bar, 20 μm. **g**, Quantification of PSMC5 expression level of (**f**). Relative PSMC5 expression level was quantified versus control scRNA+vehicle. V represents vehicle, C represents compound C. n = 3 independent experiments. Values represent means ± SEM (****p < 0.0001; ns: not significant; two-way ANOVA followed by Bonferroni test). **h**, Quantification of average neurite length of (**f**). n = 3 independent experiments. V represents vehicle, C represents compound C. Values represent means ± SEM (*p < 0.05; ***p < 0.001; ns: not significant; two-way ANOVA followed by Bonferroni test). **i**, Immunoblot shows PSMC5 and AMPKα1 expression 48 h after transfection with empty vector, WT, phospho-dead (S120A) and phospho-mimetic (S120D) PSMC5 plasmid DNA in F11-DRG cell lines. **j,** Quantification of PSMC5 and AMPKα1 expression levels of (**f**). n = 3 independent experiments. Relative protein expression level (vs Vector) was quantified after normalization to GAPDH. Values represent means ± SEM (****p < 0.0001; ns: not significant, two-way ANOVA followed by Bonferroni test). **k**, Representative neurite outgrowth images of cultured DRG neurons 36 h after electroporation with empty vector, PSMC5 WT and phospho-mimetic (S120D) plasmid. Cells were immunostained with anti-PSMC5, anti-GFP and counterstained with DAPI. Scale bar, 200 μm. **k**(i-iii), scale bar, 20 μm. **l**, Quantification of PSMC5 expression level of (**k**). n = 3 independent experiments. Relative PSMC5 expression level was quantified versus empty vector. Values represent means ± SEM (***p < 0.001; ****p < 0.0001; one-way ANOVA followed by Bonferroni test). **m**, Quantification of average neurite length of (**k**). n = 3 independent experiments. Values represent means ± SEM (****p < 0.0001; ns: not significant, one-way ANOVA followed by Bonferroni test).

It has been shown that PSMC5 activation in neurons depends upon phosphorylation at Serine 120 (S120) by Ca^2+^/calmodulin-dependent protein kinase II α (CaMKIIα)^30^, whose activity is required for signalling following nerve injury ^56^. Therefore, we tested whether PSMC5 S120 phosphorylation would affect AMPKα1 protein expression levels and DRG neurite outgrowth. To this end, we transfected F11 DRG cells with full length WT PSMC5, with PSMC5 phospho-mutant S120A, or PSMC5 phospho-mimetic S120D respectively. Immunoblotting revealed that PSMC5 S120D only significantly reduced AMPKα1 protein level, suggesting that an active PSMC5 is required to downregulate AMPKα1 (Fig. 5i, j). Likely, in these culture conditions PSMC5 remains unphosphorylated since overexpression of WT or PSMC5 S120A do not have noticeable effects upon AMPKα1 expression (Fig. 5i, j). This led to the prediction that PSMC5 S120D would promote DRG neurite outgrowth. Indeed, when we electroporated DRG neurons with PSMC5 WT or S120D, we found that cell expressing PSMC5 S120D but not WT, displayed enhanced neurite outgrowth (Fig. 5k-m). Unfortunately, endogenous PSMC5 phosphorylation cannot be measured since p-specific antibodies are not available.

Finally, we investigated whether AMPKα1 protein expression requires the activation of CaMKIIα after SNA, since active p-CAMKIIα is in turn responsible to phosphorylate and activate PSMC5. Adult mice were treated by intraperitoneal injection with the CaMKII inhibitor KN-93(12.5 mg/kg) that prevents the activation of CaMKII by calcium/calmodulin thereby reducing the autophosphorylation of CAMKIIα or with an inactive analogue of KN-93, KN-92(12.5 mg/kg). Sciatic DRGs were collected 24 h after SNA. Immunoblot showed up-regulation of phosphorylated CaMKIIα following SNA and that inhibition of the activating auto-phosphorylation of CaMKIIα with KN-93 blocked SNA-dependent reduction of AMPKα1 (Supplementary Fig. 6c, d), suggesting that CaMKIIα might be required for AMPKα1 protein expression after nerve injury. However, CaMKIIα phosphorylation did not change following DCA (Supplementary Fig. 6e, f).

### AMPKα1 conditional deletion in sensory neurons enhances key regenerative signalling molecules and promotes axonal regeneration after SCI

Initial proteomics and RNAseq data as well as the following experimental evidence so far suggested that AMPK might be play a vital role in controlling regenerative signalling. Therefore, we asked whether AMPKα1 deletion would enhance multiple key signalling pathways involved in axonal regeneration of sensory neurons (galanin, Lgals, arginase 1, p53, p21, Sprr1a, IL6, HIF1 and others), including the ones predicted by our protein-protein-interaction and functional gene network analysis (BDNF, Jun, insulin signlling-IGF1, Myc, Fos, Atf, see Fig. 1e). To investigate whether the expression of these genes would depend upon AMPKα1 expression in DRG neurons after SCI, we performed quantitative RT-PCR 24 hours after spinal cord injury and 5 weeks following AAV-cre-GFP mediated conditional deletion of AMPKα1 by injecting the cre-GFP or control GFP virus in the sciatic nerve of AMPKα1 floxed mice (Supplementary Fig. 7a, b). Data analysis revealed that conditionally deleted DRG display a significant increased expression of a number of key genes belonging to regenerative signalling pathways including Jun, p53, BDNF, Atf3, arginase 1, Fos, Myc and IGF-1, while it had no effect upon HIF1, Cxcl12 and HDAC5 (Fig. 6a). As expected AMPKα1 conditional deletion in DRG neurons led to changes in phosphorylation of well-defined protein targets including p-acetyl–coenzyme A carboxylase (pACC) and pERK (Fig. 6b-d). Thus, these data suggest that AMPKα1 deletion induces multiple regenerative signalling pathways.

**Figure 6.**
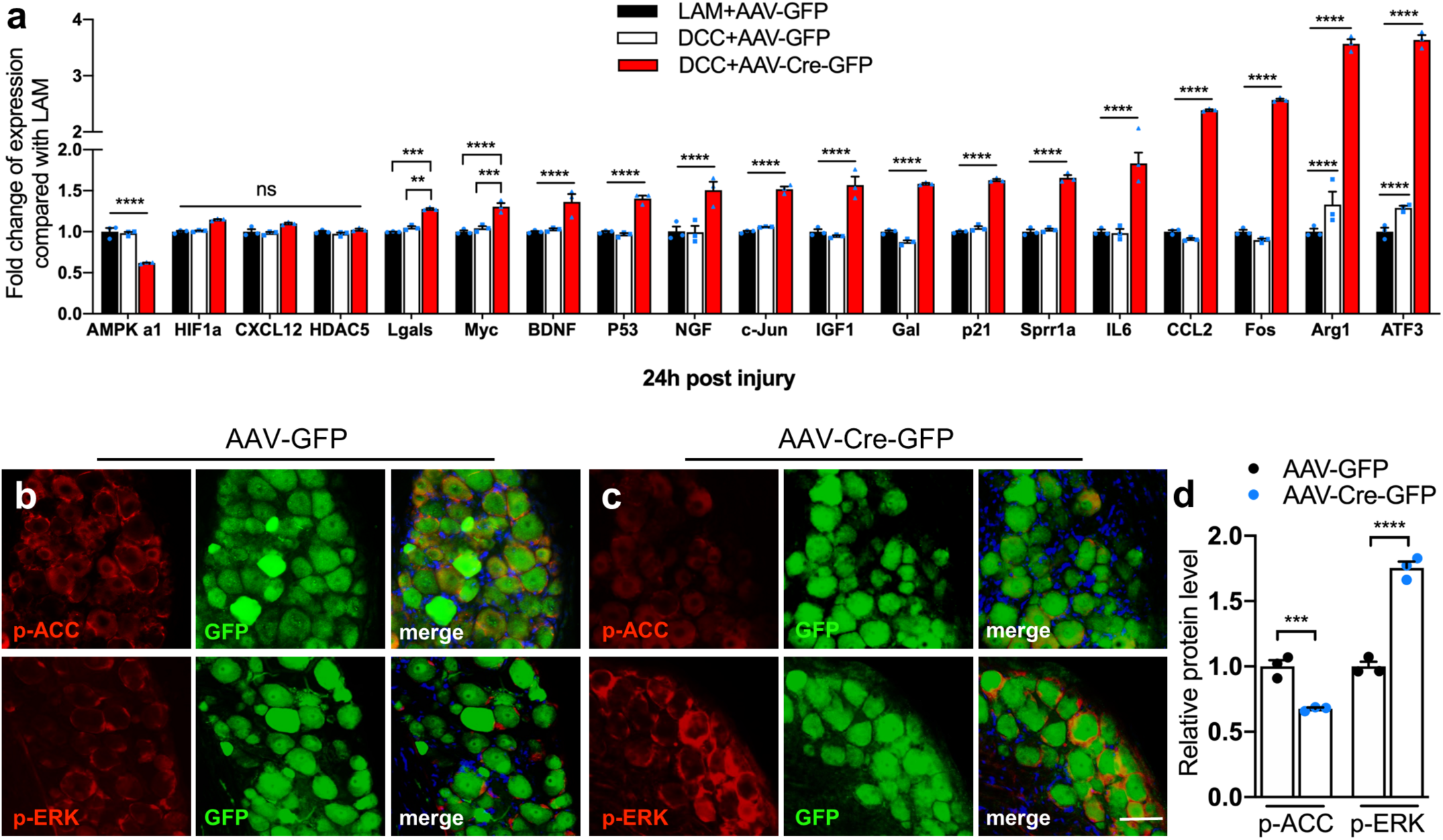
AMPKα1 regulates the expression of multiple injury-induced RAGs. **a**, Bar graphs showing quantitative RT-PCR analysis of the expression of selected RAGs in sciatic DRGs 24 h following SCI. AAV-GFP and AAV-Cre-GFP viruses were injected into the sciatic nerves 4 weeks before injury. n = 3 mice each group. The relative gene expression level was quantified versus Lam+AAV-GFP. Values represent means ± SEM (**p < 0.01; ***p < 0.001; ****p < 0.0001; ns: not significant; two-way ANOVA followed by Tukey test). **b**, **c**, Representative immunofluorescence images of p-ACC and p-ERK after conditional deletion of AMPKα1 in sciatic DRG neurons. DRG sections were immunostained for p-ACC, p-ERK together with GFP and DAPI. Scale bar, 50 μm. (d) Quantification of p-ACC and p-ERK expression levels of (**b**, **c**). n = 3 mice each group. The relative protein expression level was quantified versus AAV-GFP. Values represent means ± SEM (****p < 0.0001; two-way ANOVA followed by Bonferroni test).

Next, we investigated whether conditional deletion of AMPKα1 in the DRG would enhance axonal growth across the inhibitory spinal cord environment. To this end, we injected the sciatic nerve of AMPKα1 floxed adult mice bilaterally with an AAV-cre-GFP or an AAV-GFP control virus 4 weeks prior to a T9 spinal cord dorsal column crush to delete AMPKα1 in sciatic DRG neurons (Supplementary Fig. 7a, b). Seven days before sacrificing the animals, at day 28 post-SCI, the axonal tracer dextran was injected in the sciatic nerve to monitor axonal growth, showing co-expression in GFP positive transduced DRG (Supplementary Fig. 8a, c). Data analysis revealed that AMPKα1 deletion leads to significant labelling of traced sensory axons into and past the lesion site (Fig. 7a-c and Supplementary Fig. 8d). Interestingly, conditional deletion of AMPKα2 did not affect axonal growth, suggesting that AMPKα1 is specifically implicated in the growth of sensory axons following SCI (Fig. 7a-c).

**Figure 7.**
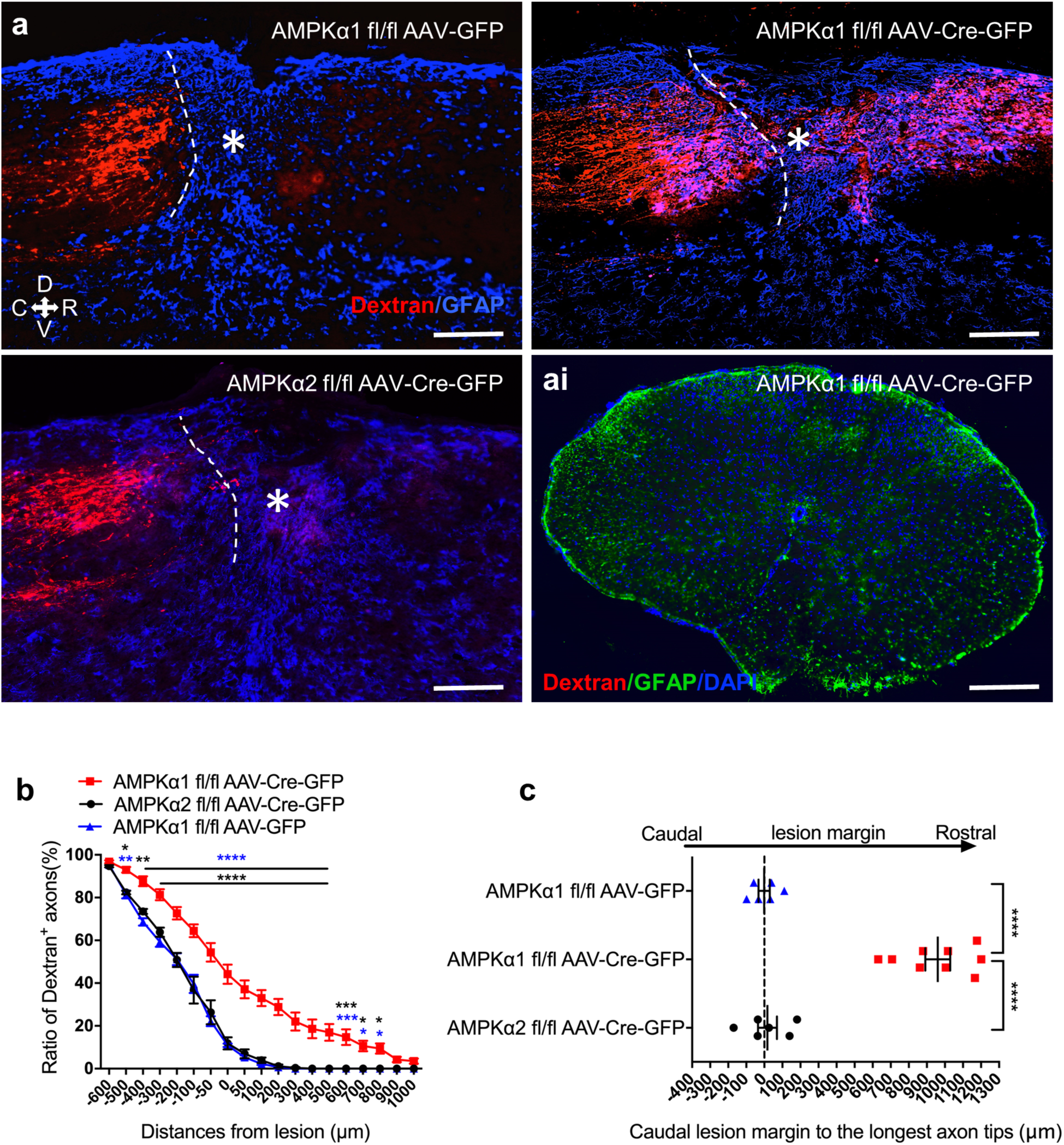
AMPKα1 deletion promotes axonal growth after SCI. **a**, Representative images of longitudinal spinal cord sections 5 weeks after SCI. Dorsal column axons are labeled by sciatic nerve injected Dextran. Asterisks indicate the lesion epicenter. Dotted lines indicate the caudal margin of the lesion site. D; dorsal; V; ventral, C; caudal; R; rostral. Scale bar; 250 μm. Figure **ai**, fluorescence image of immunostaining for GFAP with Dextran and DAPI of a spinal cord coronal section from AMPKα1^fl/fl^ mice injected with AAV-Cre-GFP. Shown is the absence of spared dextran^+^ axons at 8 mm rostral to the lesion site. Scale bar, 500 μm. **b**, **c**, Quantification of dextran positive axons. Data is expressed as ratio of dextran^+^ axons at each distance versus dextran^+^ axons at −700 μm from the lesion site (**b**) or distance from the caudal margin of the lesion to the last dextran^+^ axon tip (**c**). In **b**: blue asterisk indicates AMPKα1^fl/fl^ AAV-Cre-GFP versus AMPKα1 ^fl/fl^ AAV-GFP; black asterisk indicates AMPKα1 ^fl/fl^ AAV-Cre-GFP versus AMPKα2 ^fl/fl^ AAV-Cre-GFP. Values represent means ± SEM (*p < 0.05; **p < 0.01; ***p < 0.001; ****p < 0.0001; two-way ANOVA followed by Tukey test; AMPKα1 ^fl/fl^ AAV-Cre-GFP n = 9 mice; AMPKα1 ^fl/fl^ AAV-GFP n = 6 mice; AMPKα2 ^fl/fl^ AAV-Cre-GFP n = 6 mice). In **c**: Values represent means ± SEM (****p < 0.0001; two-way ANOVA followed by Tukey test; AMPKα1 ^fl/fl^ AAV-Cre-GFP n = 9 mice; AMPKα1 ^fl/fl^ AAV-GFP n = 6 mice; AMPKα2 ^fl/fl^ AAV-Cre-GFP n = 6 mice).

## Discussion

The differential regenerative response following an injury to the regeneration-competent peripheral versus the regeneration-incompetent central branches of dorsal root ganglia (including outside and inside the spinal dorsal columns) has been long investigated as a model to identify regenerative signalling mechanisms. Whether this dichotomic response to injury underlines differential molecular signatures in the axoplasm of central versus peripheral axons subsequent to an injury remains unexplored.

Here, we performed sciatic DRG axoplasm proteomics following a regeneration-incompetent spinal versus a regeneration-competent sciatic nerve injury in order to identify differential molecular signatures between the two. Although the sciatic nerve includes motor axons while the dorsal roots do not, the contingent of motor fibres is minimal accounting for only 6% of the total pool of fibres ^57^, likely only affecting a small minority of our axoplasmic proteins.

Gene ontology analysis showed that differentially expressed proteins in axoplasm after SNA mainly contribute to regulation of ATP metabolism and adenyl ribonucleotide signalling, transcription and translation, cell adhesion and wound healing. Differentially expressed proteins after DCA are mostly involved in translation and mitochondrial function. A number of classic regeneration-associated proteins were found upregulated in the axoplasm following sciatic injury only. GO, KEGG and combined pathway analysis found that SNA enriched proteins mainly involved in regulation of actin cytoskeleton, metabolic pathways, insulin signalling and several other regenerative pathways. However, following DCA, enriched genes and proteins were mainly involved in amino acid metabolism. Together, our data suggested that peripheral and central branch injuries lead to a distinct response both in the axoplasm and DRG. The analysis of RNAseq and proteomics data allowed us to hypothesize that AMPK might be potentially involved in the differential molecular response between the central regeneration-incompetent versus the peripheral regeneration-competent axonal lesion.

In fact, we found enrichment for AMPK regulatory mechanisms including ATP and adenyl ribonucleotide signalling, for AMPK-dependent classical metabolic pathways such as carbohydrate, amino acid metabolism ^39,47,49,50^, and for adipocytokine signalling ^51,52^. Additionally, enrichment of several regenerative pathways such as insulin, mTOR and JAK-STAT signalling, which are regulated and regulate AMPK ^26–28^ provided further evidence for a potential regulatory role for AMPK in sensory axon regeneration.

The AMPK enzymatic complex is a heterotrimer with α catalytic subunit α1 or α2 and two regulatory subunits, β (β1 and β2) and γ (γ1, γ2 and γ3) ^58^. AMPK is a key energy sensor in cellular metabolism responding to stress signalling by enhancing catabolism, fatty acid production and glucose transport at the expenses of protein and fatty acid synthesis ^59^. However, the enzymatic complex expressing AMPKα1, which we found to play a role in axonal regeneration, preferentially regulates cellular signalling rather than metabolism only ^59^. AMPK is typically regulated by phosphorylation on Thr 172 in the subunit α1 in response to ADP via the kinase LKB1 by an increase in ADP:ATP, while it is dephosphorylated by PP2A and C ^59^. However, phosphorylation of AMPK does not seem to be a major event in the axoplasm after nerve injury as suggested by lack of differentially expressed AMPK phospho-peptides after nerve injury in a sciatic nerve phosho-proteomics study in rats ^60^.

Here we show a novel mechanism of regulation of AMPK that likely depends upon PSMC5-dependent proteasomal degradation. The proteasome that is mostly exclusively used as 26S proteasome contains a catalytic core subunit (CP, also known as 20S subunit) and one or two regulatory subunits (RP, also known as 19S subunit), which serves as a proteasome activator and recognises the ubiquitinated proteins for translocation into the catalytic subunit for degradation ^61,62^. In order to clarify the mechanisms that control AMPKα expression upon SNA, we performed AMPK immunoprecipitation followed by mass spectrometry. Interestingly, we found a number of the 26S proteasomal subunits showed increased interaction with AMPKα following SNA, suggesting that the proteasome may contribute to AMPKα protein expression. Indeed, *in vivo* and *in vitro* administration of the proteasome inhibitor Bortezomib showed increased AMPKα1 expression, including after nerve injury, allowing the conclusion that the reduction of AMPKα1 following SNA includes proteasome-dependent mechanisms. Since protein network analysis identified PSMC5 as key connection node between AMPKα and the 26S proteasome, we tested whether PSMC5 would form a direct interaction with AMPKα. Indeed, AMPKα IP followed by PSMC5 immunoblot showed that these two proteins interact. PSCM5 has been reported to be a key 19S regulatory subunit for proteasome activity by modulating proteasome assembly ^55^. Accordingly, we found that silencing of PSMC5 or overexpression of PSMC5 S120D promoted or blocked AMPKα1 expression with decreased or increased neurite outgrowth respectively. Additionally, we established that reduction of CaMKIIα phosphorylation is followed by SNA-dependent decline of AMPKα1 expression in DRG and that overexpression of phospho-mimetic S120D PSMC5, which has been shown to be phosphorylated and activated by CaMKIIα ^63^, reduces AMPKα1 expression and promotes DRG regenerative growth. It is interesting to underline how AMPKα2 expression is not affected by either a peripheral sciatic or central spinal injury. Additionally, AMPKα2 expression is mainly nuclear while AMPKα1 localises to the cytoplasm, suggesting very different roles in DRG neurons. Indeed, both AMPKα1 and 2 subunits co-immunoprecipitate with PSMC5, while only AMPKα1 protein level decreases after SNA and is affected by proteasomal inhibition. This suggests that PSMC5 interaction with nuclear AMPKα2 might underline a proteasomal-independent role of PSMC5 including the regulation of transcriptional-dependent signalling as previously suggested for both PSMC5 and AMPKα2 ^64–66^. These likely regulate mechanisms that are independent from sensory axonal regeneration since AMPKα2 deletion in DRG neurons does not affect their regenerative ability.

Our discovery that AMPKα1 conditional deletion in sciatic DRGs induces the expression of multiple regeneration associated genes belonging to multiple signalling pathways following spinal cord injury expands upon published findings showing that AMPK signalling has profound effects on gene expression in other cell types and model systems. Activation of AMPK in cell types other than neurons has been shown to inhibit the expression and activity of several regenerative TFs including STAT3, CREB, c-jun and the histone acetyltransferase p300 involved in the regeneration programme ^67–71^. Another study in cultured neurons found that AMPK can directly phosphorylate kinesin light chain 2 (KLC2) and inhibit neurite growth through prevention of PI3 K localization at the axonal tip ^72^. Indeed, AMPK inhibition protects against autophagy and enhances mTOR signalling via the de-phosphorylation of tuberous sclerosis 2 (TSC2), both beneficial for neuronal metabolism ^15,73^ and potentially affecting axon regeneration. Additionally, GSK3β, whose downregulation stimulates axonal regeneration of sensory neurons ^74^, inhibits the mTOR pathway by phosphorylating TSC2 in a manner dependent on AMPK-priming phosphorylation ^73^. Interestingly, IGF1 signalling, recently shown to promote axonal regeneration by others and us ^75,76^, is inhibited by AMPK activation ^77^. Finally, deletion of PTEN, a powerful axonal regenerative stimulus ^15^, inhibits AMPK signalling ^78^, while AMPK activates PTEN ^79^, confirming our data suggesting AMPK as a crucial regulator in the control of the axonal regeneration programme.

Indeed, conditional neuronal deletion of AMPKα1 promotes axonal growth of sensory axons after SCI.

In summary, we provide novel axoplasmic expression protein profiles represent a resource for the community interested in understanding the injury-dependent molecular response of peripheral versus central DRG projections. Next, we show the evidence that the axoplasmic protein expression signatures of central and peripheral projecting sciatic DRG axons subsequent to an injury are unique. Lastly and more importantly, we identify a mechanism for AMPK regulation in DRG neurons and propose AMPK as a novel inhibitor of sensory axonal growth ability following a spinal cord injury whose deletion promotes robust axonal growth.

## Acknowledgments

We would like to thank start-up funds from the Division of Brain Sciences, Imperial College London (SDG), and the Hertie Foundation for financial support (SDG); the ISRT (Elisabeth Serger and Eilidh Maclachlan); Wings for Life (SDG); the DFG, the MRC (SDG), and Rosetrees Trust (SDG). The research was supported by the National Institute for Health Research (NIHR) Imperial Biomedical Research Centre (SDG). The views expressed are those of the author(s) and not necessarily those of the NHS, the NIHR or the Department of Health. Q Exactive Plus mass spectrometer was funded by DFG - Deutsche Forschungsgemeinschaft INST 247/766-1 FUGG. We would also like to thank Monica Sousa and Mike Fainzilber for critical discussing the data and the manuscript.

## SUPPLEMENTARY MATERIAL

### METHODS

#### Animals

All animal procedures were performed in accordance to The Animal Welfare Act and the guidelines of the University of Tuebingen. Three mouse germ-lines were used for this study: C57BL6/J (Charles River Laboratories), Prkaa1^fl^ (Stock No: 014141) and Prkaa2^fl^ (Stock No: 014142) mice (purchased from The Jackson Laboratory). An equal number of male and female animals was used for all the experiments.

#### AAVs

AAV-GFP and AAV-Cre-GFP were purchased from SignaGen Laboratories.

#### Chemical reagents

Compound C was purchased from Sigma (866405-64-3). Bortezomib was obtained from Selleckchem (PS-341). KN-93 and KN-92 phosphate were purchased from MedChem Express (HY-15465B, HY-15517A).

#### Plasmids

The PSMC5 WT, phospho-dead(S120A) and phospho-mimetic (S120D) were obtained from Prof. Gentry Patrick, University of California, San Diego. The AMPKα1 plasmid (HCG11488-ACG) was purchased from Sino Biological.

#### siRNA

The control siRNA (sc-37007) and PSMC5 siRNA (SC-76604) were purchased from Santa Cruz.

#### Axoplasm preparation for proteomics

6-8-week old mice underwent bilateral sciatic nerve axotomy or sham injury. The axotomy injury was performed at about 1.5 cm distally to the sciatic DRG. 24 h later, animals were sacrificed and the proximal nerve segments were collected in 500 µl 0.2X PBS on ice and then processed for axoplasm extraction as described previously ^80^. Briefly, nerve fascicles were separated carefully by using fine forceps, then once they became cloudy, they were transferred to a fresh eppendorf tube containing 500 µl 0.2X PBS. Following 2 h incubation at room temperature, the fascicles were washed 3 times using the same solution by transferring them from one eppendorf tube to another following 5 min shaking every time. Thereafter, the fluid was removed by placing the fascicles into a new eppendorf tube, then the axoplasm was extracted by using 300 µl 1X PBS for 30 min incubation at room temperature with subsequent centrifugation at 10,000×g for 10 min at 4°C. Protease (Roche, 04693116001) and phosphatase inhibitors (Roche, 04906837001) were added into all solutions used in the purification procedure. Finally, purified axoplasm was concentrated by centrifugation at 4,000×g for 30 min at 4°C using the Amicon Pro Purification System (Millipore, ACS500312), afterwards, 500 µl denaturation buffer (6M urea, 2M thiourea in 10 mM Tris pH 8.0) was added into the same concentrator for buffer exchange. After centrifuging at 4,000×g for 2 h at 4°C, the total volume was concentrated to about 30 µl for future LC-MS/MS analyses.

Sciatic DRG dorsal roots were collected at 24 h following Laminectomy or Dorsal column axotomy. Central branches axoplasm purification was performed following the same protocol as for the peripheral branches.

15 mice were used for each group and biological triplicated were performed with each condition.

#### AMPKα immunoprecipitation for mass spectrometry

Bilateral sciatic nerve axotomy or sham injuries were performed in 8 weeks old mice. 6 h later, sciatic DRG neurons were harvested and lysed in IP lysis buffer (25 mM Tris-HCl pH 7.4, 150 mM NaCl, 1% NP-40, 1 mM EDTA, 5% glycerol) containing a cocktail of protease and phosphatase inhibitors on ice for 30 min. After centrifuging at 12,000×g for 10 min at 4°C, protein concentration was measured by BCA protein assay kit (Thermo Fisher Scientific, 23227).

1.2 mg protein per group was used for the AMPKα immunoprecipitation experiment. 5 µg AMPKα antibody (Abcam, ab80039) was added into each sample for overnight incubation at 4°C on a rotary device. Normal rabbit IgG immunoprecipitation was used as negative control. Each condition (Sham or SNA) has their own IgG control. On the second day, 50 µl protein A Dynabeads (Thermo Fisher Scientific, 10001D) was washed 2 times with 500 µl IP lysis buffer and incubated with the protein and AMPKα antibody mixed solution for 1 h at 4°C. Thereafter, the nonspecific proteins were separated by the Magna GrIP Rack (Millipore, 20-400). The Dynabeads complex was washed 3 times with 1 ml IP lysis buffer. Next, Dynabeads were resuspended in 30 µl 1X NuPAGE LDS loading buffer (Thermo Fisher Scientific, NP0007) containing reducing agent and mixed gently. Next the samples were boiled 10 min at 70°C, the LDS loading buffer was separated from the beads mixture and collected for mass spectrometry analyses. All samples were prepared in biological duplicates.

#### Mass Spectrometry Sample Preparation and Measurement

Samples were boiled at 70°C for 10 min in 1x NuPAGE LDS Sample Buffer (Thermo Fisher Scientific, NP0007) containing 100 mM DTT and separated on a 10% NuPAGE Bis-Tris gel (Thermo Fisher Scientific, NP0301BOX) for 10 or 20 min at 180 V in MES running buffer (Thermo Fisher Scientific, NP000202). After fixation in 7% acetic acid containing 40% methanol and subsequent staining for 30 min using Colloidal Blue staining kit (Thermo Fisher Scientific, LC6025), protein lane was excised from the gel (for the axoplasm proteome analysis samples was separated in six slices), chopped and destained (50% ethanol in 25 mM NH_4_HCO_3_) for 15 min rotating at room temperature and dehydrated for 10 min rotating in 100% acetonitrile. Vacuum dried samples were rehydrated and reduced for 60 min in reduction buffer (10 mM DTT in 50 mM NH_4_HCO_3_ pH 8.0) at 56°C and subsequently alkylated in 50 mM iodoacetamide in 50 mM NH_4_HCO_3_ pH 8.0 for 45 min at room temperature in the dark. Dehydrated and vacuum dried samples were trypsin digested (1 µg trypsin/sample in 50 mM Triethylammonium bicarbonate buffer pH 8.0) at 37°C overnight. Stepwise peptide extraction was done as follows: twice extraction solution (30% acetonitrile) and 100% acetonitrile for 15 min at 25°C shaking at 1,400 rpm. Reductive methylation for quantification was performed as previously described^81^. After purification and desalted using C18 stage tips 3.5 µl peptides were loaded and separated on C18 column (New Objective) with 75 µm inner diameter self-packed with 1.9 µm Reprosil beads (Dr. Maisch) which was mounted to an EasyLC1000 HPLC (Thermo Fisher Scientific, LC120). Reversed-phase chromatography gradient (Buffer A: 0.1% formic acid, Buffer B: 80% acetonitrile and 0.1% formic acid, Gradient: 0-67 min 0-22% Buffer B, 67-88 min 22-40% Buffer B, 89-92 min 40-95% Buffer B) was applied and peptides eluted and directly sprayed into a Q Exactive Plus mass spectrometer (Thermo Fisher Scientific, IQLAAEGAAPFALGMBDK) operating in positive scan mode with a full scan resolution of 70,000; AGC target 3×10^6^; max IT = 20 ms; Scan range 300 - 1650 m/z and a Top10 MSMS method. Normalized collision energy was set to 25 and MSMS scan mode operated with resolution of 17,000; AGC target 1×10^5^; max IT = 120 ms.

Database search was performed using MaxQuant Version 1.5.2.8^82^ against Mus Musculus Ensembl database (release-81; 3rd July 2015; 53,819 entries) for axoplasm proteome analysis and Mouse Uniprot database (downloaded 8th of January 2015; 83,429 entries) for AMPK immunoprecipitation analysis, with Trypsin/P as digestion enzyme allowing 2 missed cleavages. As settings the following was applied: variable modification: Acetyl (Protein N-term); Oxidation (M), fixed modifications: Carbamidomethyl (C), FDR of 1% on peptide and protein level was applied.

As light label: DimethylLys0 and DimethylNter0 and heavy label: DimethylLys4 and DimethylNter4 were set with max. 3 labelled amino acids.

Proteins with at least two unique peptides were considered as identified. Proteins matching reverse database or common contamination list as well as proteins with peptides only identified by peptides with modification were filtered out. For quantification we used MaxQuant default settings with a minimum ratio count of 2, i.e. area under the curve of min. Two unique peptides per protein were used to calculate the ratio between heavy and light. Ratio distribution was normalized by MaxQuant software in such a way that the median ratio is located close to 1. This corrects for mixing errors after dimethyl-labelling procedure.

Package “biomaRt” was used to convert Ensemble identifiers into gene names.

#### Data availability

The axoplasmic proteomics and AMPK IP followed by mass spectrometry data have been deposited to the ProteomeXchange Consortium via the PRIDE^83^ partner repository with the datasets identifiers as follows:

Axoplasmic proteomics: PXD013297

Reviewer account details: Username. Password: 91HtQLEG

AMPK IP MS: PXD013318

Reviewer account details: Username. Password: O8jGsu8C

#### Axoplasm proteome analysis

Protein abundance was quantified as a log2 ratios of Heavy (case) and Light (control) labelled samples in three replicates. Peptide measurements mapped to the same protein were merged by taking the median value. We removed less reliable data by removing measurements in only one replicate. The remaining values were combined across the three replicates by taking the mean. To determine which proteins were significantly detected above technical noise we used signal deconvolution method as implemented in R mixtools ^84^, to extract the distribution of the noise signal. We assume that the signal observed from the quantification experiment can be modelled as the combination of changes due to technical noise and actual protein levels.

Technical noise is normally distributed and centred on zero, while biological signal is normally distributed along the extreme edges of the overall distribution. After modelling the noise distribution, the remaining data follows two distributions describing the proteins that are up and down regulated in cases relative to controls. To quantify the confidence of detection of all peptides relative to the null, we performed a one-tailed *t*-test of each detected peptide relative to the null and adjusted the p-values using Benjamini-Hochberg multiple testing correction.

Heat map of the log2 ratio of significantly (FDR < 0.05) differentially expressed proteins identified by mass spectrometry in the axoplasmic extract was created using function “heatmap.2”. Comparisons include peripheral nerve after sciatic nerve axotomy (SNA) vs sham (control injury) and central branches after dorsal column axotomy (DCA) vs Lam (control injury). 389 protein groups with a log2 value in at least one comparison were plotted in a heatmap. Red and Blue indicates up- and down-regulated proteins, respectively. In white, not identified/quantified proteins are represented. Hierarchical clustering was computed on rows using “hclust” function. Gene ontology (GO) and KEGG Pathway analysis were performed using DAVID (https://david.ncifcrf.gov/) setting all the expressed genes/proteins in our dataset as background.

Transcriptomic and proteomic data have been integrated following published approaches ^41,42^. Briefly, merged the datasets considering a union of the proteomic and transcriptomic data sets to create a reference data set, which was then used for a joint pathway analysis.

Protein-protein interaction network was visualized using Cytoscape (http://www.cytoscape.org/). The AMPK interaction network was produced by generating a protein-protein interaction network using String (https://string-db.org/), integrating all proteins and genes assigned to significantly enriched pathways in the combined (Proteomics and RNAseq) KEGG analysis (p < 0.1). Genes were jointly organized in a circular layout according to their KEGG affiliation by using Cytoscape.

#### AMPKα immunoprecipitation analysis

For the AMPK immune-precipitation experiments, proteins were quantified in two replicates as forward and reverse log2 ratios of heavy and light chains. In forward comparison, case (Sham or SNA AMPKα IP) was labelled with heavy isotopes(H), IgG in each condition was labelled with Iight isotopes (L), the labelling was in the opposite way in reverse comparison. Since one of the ratios was taken with respect to controls and the other with respect to cases, they measured the same variance but the value of proteins log2 ratios are in reverse. Therefore, the absolute value of the reverse ratio was taken to match the forward observation. The average of the ratios was subsequently used to summarise the signal. In this experiment, we were interested in proteins which were significantly pulled down by AMPKα in both Sham and SNA relative to their own IgG controls. Mixtools were used to identify the proteins significantly detected above noise and log2 ratios were used to identify the peptides enrichment relative to controls. To that end we performed a one-tailed *t*-test, identifying proteins that were significantly more strongly enriched than experimental noise. We were also interested in proteins significantly more strongly interact with AMPKα in either SNA or Sham. To this end we combined statistically verified AMPKα interactors from both IP experiments (sham vs IgG and SNA vs IgG) using Fisher’s method ^85^. This method is well suited for combining information from related studies, thus reinforcing observations that follow the same trend in both experiments.

#### Immunoblotting

Bilateral sciatic DRG neurons were collected in PBS and lysed in RIPA buffer at different time points after sciatic nerve axotomy. All the buffers used in the above steps contain protease and phosphatase inhibitors. Proteins sample from all conditions were loaded on 10% SDS-PAGE gel and transferred on the nitrocellulose membrane (Thermo Fisher Scientific, IB301001) using the iBlot® Gel Transfer Device (Thermo Fisher Scientific, IB1001EU). The membrane was blocked using 5% milk for 1 h at room temperature and incubated with primary antibodies diluted with 5% milk overnight at 4°C followed by HRP-conjugated Amersham ECL Rabbit IgG (GE Healthcare Life Sciences, NA934, 1:2000) or Amersham ECL Mouse IgG (GE Healthcare Life Sciences, NA931, 1:2000) on the second day.

F11 DRG cell line lysate and DRG cell lysate were obtained by washing cells 2 times with cold PBS and lysing in RIPA buffer 48 h after siRNA and plasmid transfection. Collected protein samples were processed for immunoblotting as described above.

Sciatic axoplasm and dorsal root axoplasm were collected using the standard method mentioned above. After extraction, Amicon® Ultra Centrifugal Filter concentrated axoplasm was processed for immunoblotting as described above.

1.6 mg DRG protein was used for AMPK IP for validation of candidate protein interactors, 6 mg DRG protein was used for AMPKα1 or AMPKα2 IP. After performing standard IP protocol, the IP samples were processed by standard immunoblot procedures.

The primary antibodies used were: anti-AMPKα1 (Abcam, ab32047, 1:500), anti-AMPKα2 (Abcam, ab3760, 1:500), anti-phospho-AMPKα (Cell Signaling Technology, #2535, 1:1000), anti-PSMC5 (Abcam, ab178681, 1:5000), anti-phospho-CaMKII (Cell Signaling Technology, #3361, 1:1000), anti-CaMKIIα (Thermo Fisher Scientific, MA1-048, 1:1000), anti-GAPDH (Cell Signaling Technology, #2118S, 1:10000), anti-anti-βIII Tubulin or TUJ1 (Promega, G712A, 1:10000), anti-SNAP25 (Abcam, ab5666, 1:1000), anti-STMN2 (NOVUS, NBP1-49461, 1:1000), anti-GAP43 (NOVUS, NBP1-41123SS, 1:500), anti-IGF2BP2 (St John’s Laboratory, STJ116038, 1:500), anti-Arginase-1 (Cell Signaling Technology, #9819, 1:1000), anti-CHMP3 (antibodies-online, ABIN5969355, 1:500), anti-PSMF1 (St John’s Laboratory, STJ27500, 1:500), anti-CLIC1 (St John’s Laboratory, STJ115639, 1:500), anti-S100 (Abcam, ab52642, 1:5000), anti-MPZ (Abcam, ab31851, 1:5000), anti-MOG (Abcam, ab109746, 1:1000), anti-GPHN (St John’s Laboratory, STJ111307, 1:500), anti-CRMP 1(St John’s Laboratory, STJ23226, 1:500), anti-PSMD13 (St John’s Laboratory, STJ29036, 1:500) and anti-PSMD7 (St John’s Laboratory, STJ27309, 1:500).

#### AMPK Activity Assay

Adult mice underwent bilateral sciatic nerve axotomy or sham injury. 24 h later, sciatic DRG were collected and then lysed in lysis buffer (25 mM Tris-HCl pH 7.4, 150 mM NaCl, 1% NP-40, 1 mM EDTA, 5% glycerol) containing a cocktail of protease (Roche, 04693116001) and phosphatase inhibitors (Roche, 04906837001). After AMPKα immunoprecipitation, 1 µg DRG protein was used for the activity assay by using the CycLex® AMPK Kinase Assay Kit (MBL, CY-1182) according to the manufacture’s protocol. Measurement was performed in biological triplicate.

#### Dorsal Root Ganglion (DRG) culture and electroporation

6-8 weeks old mice were used for this experiment. DRGs were dissected and collected in HBSS on ice. Collected DRGs were digested after centrifugation in a solution of Dispase II 10 mg/ml (Sigma, D4693) and Collagenase II 20 mg/ml (Sigma, 9001-12-1) in DMEM GlutaMAX Supplement at 37°C for 35 min. Then DRG tissues were transferred in DRG media (10% FBS, 2% B27 in DMEM/F12, GlutaMAX Supplement) and mechanically dissociated into cell suspension by gentle trituration with a fire-polished sigmacote coated pipette. After cell counting, DRG cells were spun down and resuspended in DRG culture media (1% Penicillin/Streptomycin, 2% B27 in DMEM/F12, GlutaMAX Supplement). 3000-5000 cells were plated per laminin pre-coated coverslip (laminin 2.4 µg/µl (1.2 mg/ml, Millipore, CC025) or myelin 1.3 µg/cm^2^). To allow neurite outgrowth, the culture plate was placed into the incubator (37°C, 5% CO_2_) for 18 - 24 h.

For the electroporation experiment, dissociated DRG cells were spun down after counting at 800 g for 5 min. During this period, the transfection solution was prepared by mixing the Lonza nucleofection solution with GFP (0.4 µg), siRNA (6 pmol) and DNA (3–4 µg) to a final volume of 20 µl for each transfection. Then the nucleofection solution was gently mixed with the cell pellet and transferred to the electroporation cuvette. After electroporation, pre-warmed DRG culture media was added into the cuvette and cells were plated on a coverslip. The culture plate was placed into the incubator (37 °C, 5% CO_2_) for 36 - 48 h.

#### DRG neurite length analysis

The average neurite length of cultured DRG cells was measured by using Neurolucida software with at least 100 cells per condition in technical and biological triplicate.

#### Immunocytochemistry

Cultured DRG cells were fixed with 4% PFA for 20 min at room temperature and washed with PBS 3 times. Then cells were treated with 0.25% TritonX-100 for 10 min followed by washing once with PBS. Next, cells were incubated with primary and Alexa Fluor secondary antibodies at room temperature for 1 h each. All coverslips were mounted with VECTASHIELD anti-fade mounting medium (VECTOR LABORATORIES, H-1000). The primary antibodies used were: anti-βIII Tubulin or TUJ1 (Promega, G712A, 1:1000), anti-GFP (Abcam, ab13970, 1:500), anti-PSMC5 (Abcam, ab178681, 1:200) and anti-AMPKα1 (Abcam, ab32047, 1:200).

#### Immunohistochemistry

Mice were anaesthetized and perfused with 4% PFA in PBS. DRGs and spinal cords were dissected and post-fixed in PFA on ice for 2 h and then cryoprotected in 30% sucrose for 72 h at 4°C. The DRGs and spinal cord were sectioned in 10 μm and 18 μm thickness separately. The slides were treated with 5% NGS and 0.3% TritonX-100 and then incubated with primary antibodies at 4°C overnight. The next day, slides were washed with TBST 3 times and then incubated with Alexa Fluor secondary antibodies for 1 h at room temperature following TBST washing. All slides were mounted with VECTASHIELD anti-fade mounting medium (VECTOR LABORATORIES, H-1000). The primary antibodies used in these studies were: anti-GFAP (Millipore, AB5804, 1:500), anti-AMPKα1 (Abcam, ab32047, 1:100), anti-AMPKα2 (Abcam, ab3760, 1:400), anti-Neurofilment 200 (Sigma, N5389, 1:400), anti-GFP (Abcam, ab13970, 1:500), anti-p-ACC (Cell Signaling Technology, 11818, 1:100), anti-pERK (Cell Signaling Technology, 4370, 1:100), anti-PARV (Sigma, P3088, 1:500), anti CGRP (Abcam, ab81887, 1:100) and anti-IB4 (Millipore, 217660, 1:100).

#### Cell culture and transfections

F11 DRG cells were cultured in DMEM GlutaMAX supplement with 10% FBS, 2 mM L-Glutamine and 1% Penicillin/Streptomycin. The day before transfection, cells were seeded into a 24-well plate to reach 60% - 80% density on the second day when transfection was performed. Transfections of the siRNA and plasmids were performed using Lipofectamine RNAiMAX (Thermo Fisher Scientific) and Lipofectamine 3000 (Thermo Fisher Scientific) respectively, according to the manufacture’s protocol. 48-72 h later cells were harvested for immunoblotting.

#### RNA isolation and reverse transcription

Bilateral sciatic DRGs were harvested in RNAlater Stabilitation Solution and total RNA was extracted according to the manufacture’s protocol (Rneasy Mini Kit, QIAGEN). RNA quality was assessed by measuring the ratio of absorbance at 260 nm and 280 nm using a NanoDrop 2000 spectrometer (Thermo Fisher Scientific). Reverse transcription was performed using SuperScript™ II Reverse Transcriptase (Thermo Fisher Scientific), according to the manufacture’s protocol.

#### Quantitative real-time PCR

qRT-PCR was performed using KAPA SYBR® FAST qPCR kit (Sigma, KK4601) in Applied Biosystems® 7500 Real-Time PCR System. The target gene expression level was normalized to the housekeeping gene GAPDH. Primers used in qPCR were as follows:

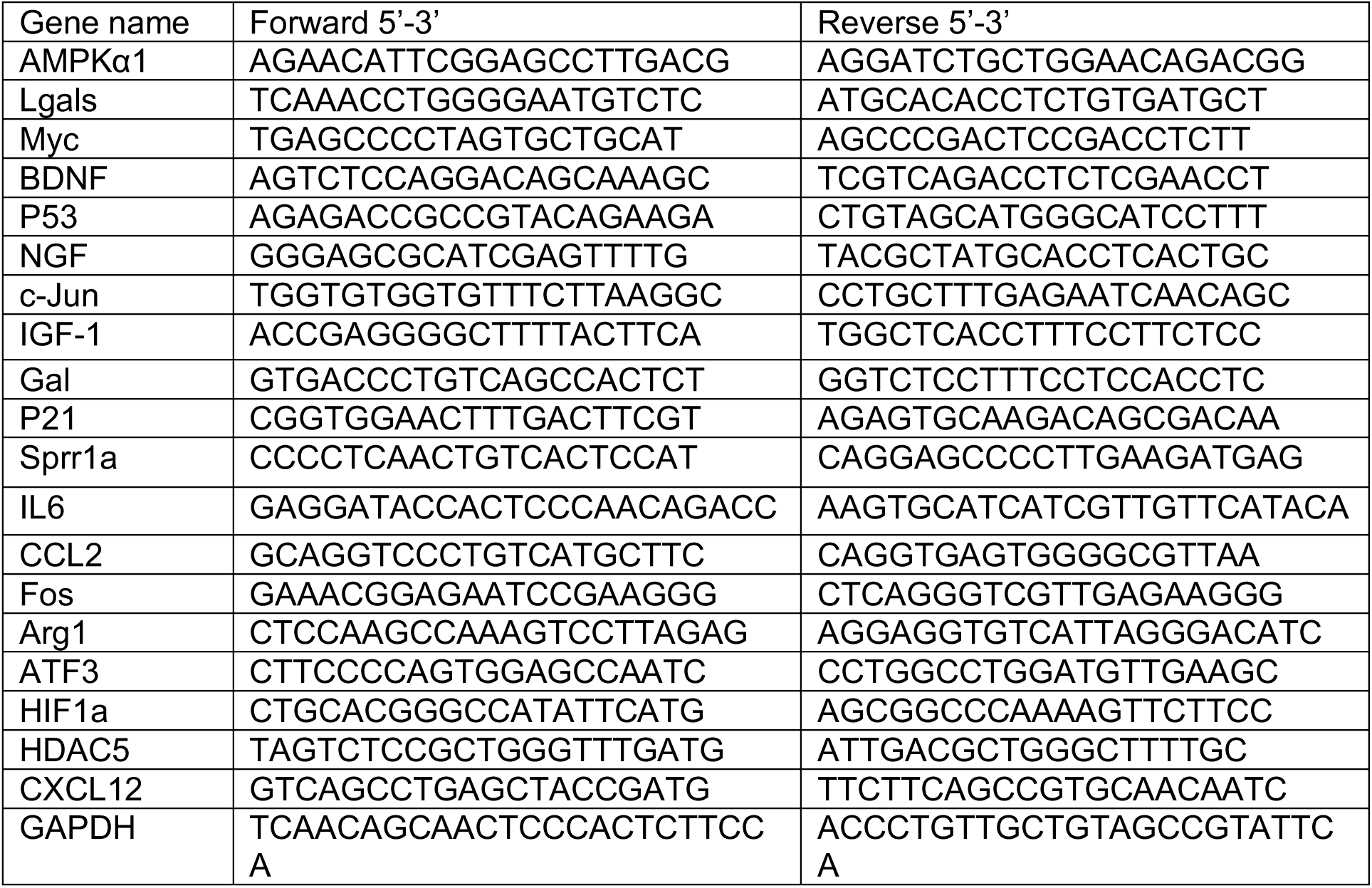

#### Peripheral nerve crush

4-6 weeks old mice were anaesthetized with xylazine (10 mg/kg of body weight) and ketamine (100 mg/kg of bodyweight). After shaving, the sciatic nerve was exposed at the middle thigh level and crushed 15 s using #2 forceps (Dumont, FST).

#### Viral injections and dorsal column crush

Four-week old AMPKα1 (prkaa1)^fl/fl^ and AMPKα2 (prkaa2)^fl/fl^ mice were anaesthetized with xylazine (10 mg/kg of body weight) and ketamine (100 mg/kg of bodyweight) and sciatic nerves were injected bilaterally with 2 µl AAV-GFP or AAV-Cre-GFP using a Hamilton syringe attached with a glass-pulled micropipette.

Four weeks after the AAV injection, a T9 dorsal column crush was performed. Specifically, a T9-T10 laminectomy was performed and it was followed by a dorsal column crush lesion to a depth of 0.5 mm for 5 s with forceps (Dumont #5, FST).

#### Dextran tracing and tissue processing of injured spinal cord

In order to retrogradely label ascending regenerative axons running into the dorsal columns, four weeks after injury (7 days before sacrificing mice), 2 µl of dextran (Thermo Fisher Scientific, D34679) was injected into the sciatic nerve bilaterally. Spinal cords were dissected 7 days after tracing following transcardiac perfusion with PBS and 4% PFA, and subsequently cryopreserved with 30% sucrose at 4°C for 72 h. Spinal cords were mounted and frozen in OCT mounting media and sectioned at 18 μm on a cryostat. Subsequent immunostainings were performed according to standard procedures. A high-resolution image was obtained at 20x magnification using the Zeiss Axioplan 2 Microscope (Axiovert 200, Zeiss Inc.).

#### Quantification of axonal regeneration

For each spinal cord after dorsal column injury, the number of fibers rostral to the lesion and their distance from the lesion epicenter and margin was analyzed in 4-6 sections per animal with a fluorescence Axioplan 2 (Zeiss) microscope and with the software Stereo-Investigator 7 (MBF bioscience). The lesion epicentre and margin (GFAP) was identified in each section at 40x magnification. The total number of labelled axons at each distance to the lesion site was normalized to the total number of labelled axons at 700 μm caudal to the lesion site and was counted in all the analyzed sections for each animal, obtaining an inter-animal comparable ratio. Sprouts and regrowing fibers were defined following the anatomical criteria reported by Steward and colleagues ^86^. Coronal section of spinal cord at 8 mm rostral to the lesion was used to ascertain that the dextran positive axons across the lesion site were regenerating axons and not spared fibers.

#### Measurement of fluorescence intensity

Images of DRG sections or cultured DRG neurons were taken at 10x or 20x magnification using Zeiss Axioplan 2 Microscope (Axiovert 200, Zeiss Inc.). Exposure time and gain were maintained constant between conditions for each fluorescence channel. Using Image J, the intensity values of each cell were normalized to the background fluorescent signal and mean values of intensities were calculated for each animal or sample. All measurements were done in triplicate and blind to the experimental group.

#### Statistical Analysis

All statistical analyses were performed using Graphpad Prism 8.0 (Graphpad Software Inc., La Jolla, CA). Unless otherwise stated, data is plotted as the mean ± SEM. All experiments were performed three times unless specified. Asterisks indicate a significant difference analyzed by one-way or two-way ANOVA with Bonferroni test, Dunnett test, Tukey test or unpaired Student’s *t*-test as indicated for normal distributions (*p < 0.05; ** p < 0.01; ***p < 0.005; ****p < 0.001). All tests performed were two-sided, and Fisher’s method was used for multiple comparisons and/or significantly different variances analysis when appropriate. All data analysis was performed blind to the experimental group.

**Supplementary Figure 1.**
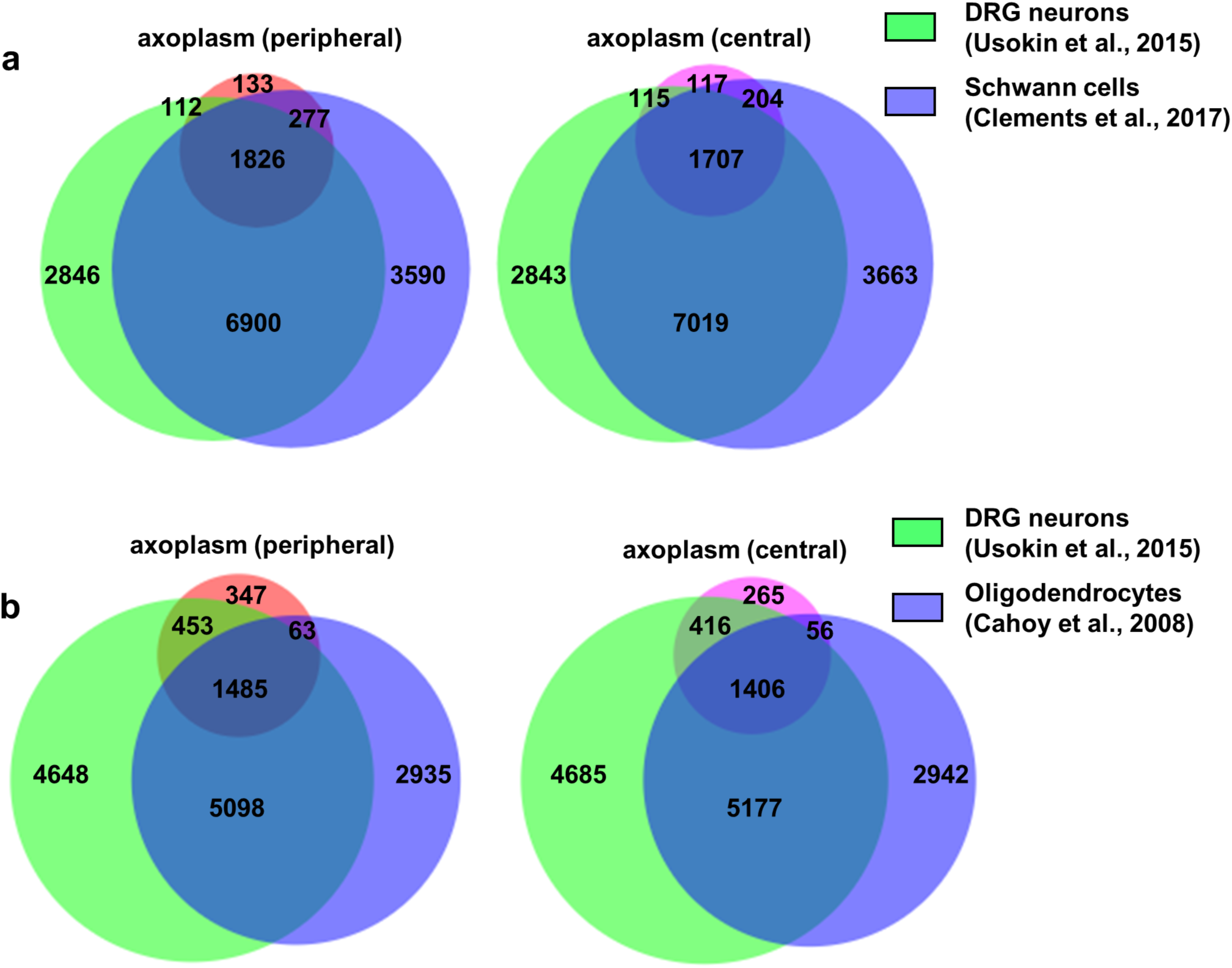
**a**, Area-proportional Venn diagram showing the overlap between the proteins identified in our axoplasm dataset and previously published DRG neuron or Schwann cell specific transcriptomic datasets. Red: peripheral axoplasm; pink: central axoplasm. **b**, Area-proportional Venn diagram showing the overlap between proteins represented in our axoplasm dataset and previously published DRG neuron or Oligodendrocyte specific transcriptomic datasets.

**Supplementary Figure 2.**
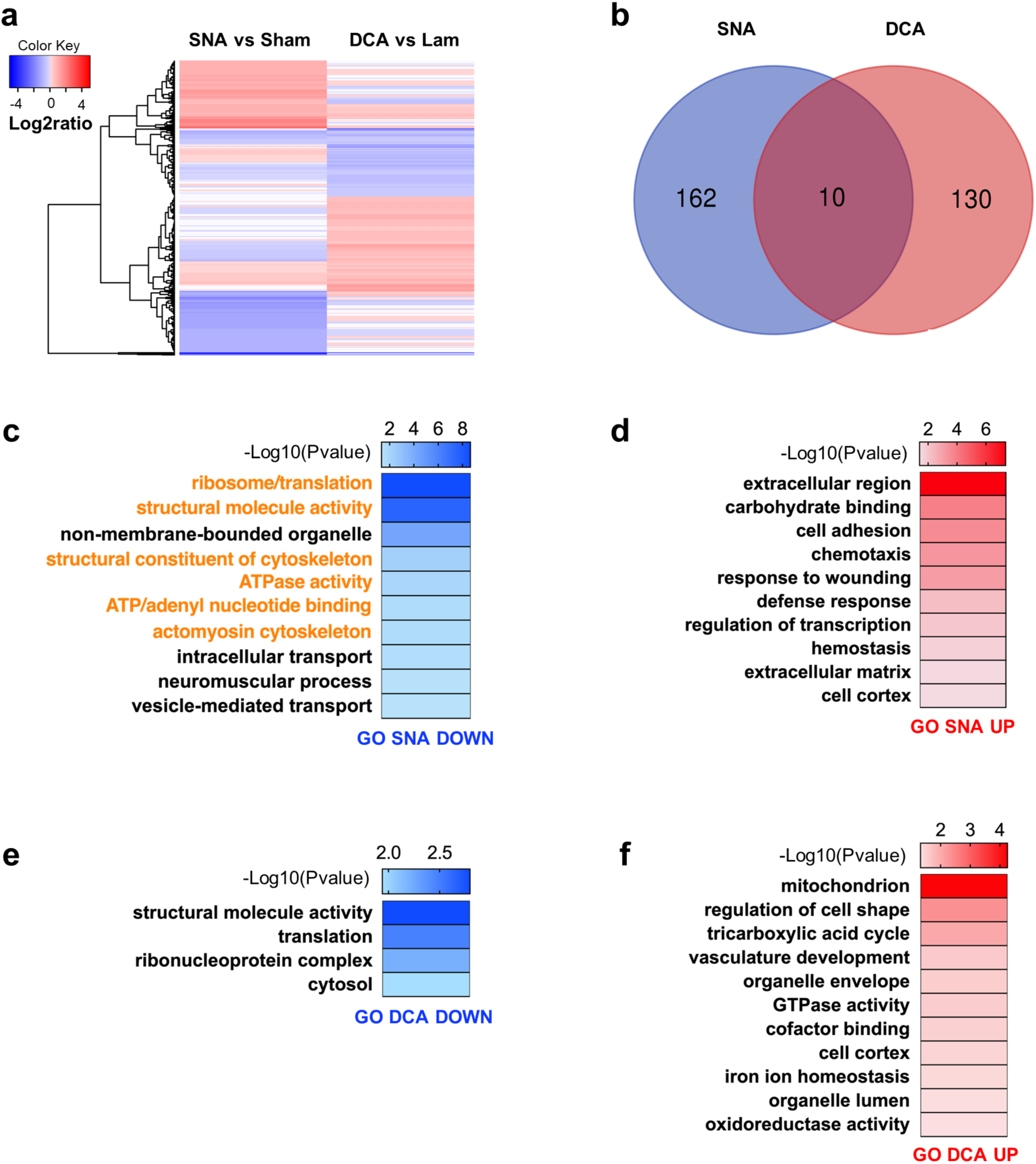
Injuries to the DRG peripheral and central axons elicit differential protein expression profiles. **a**, Heat map of the log2 ratio of differentially expressed proteins (FDR < 0.05) identified by mass spectrometry in the axoplasmic extract from peripheral and central DRG axons. Comparisons include peripheral nerve after sciatic nerve axotomy (SNA) vs sham (control injury); central branches after dorsal column axotomy (DCA) vs Lam (control injury). Red and Blue indicates up- and down-regulated proteins, respectively. **b**, Venn diagram shows the number of differentially expressed proteins following SNA and DCA (FDR < 0.05, |log2ratio| > 0.58) and how many proteins are overlapped between these two compartments after injury. **c**-**f**, Heatmap graphs show Gene ontology (GO) analysis of differentially expressed proteins following SNA and DCA. Differentially expressed proteins were selected with cut off FDR < 0.05, log2 ratio > 0.58 (Red) or log2 ratio < −0.58 (Blue). Gene ontology was performed by DAVID. Only enriched GO items with p-value < 0.05 were selected and categories that share the same protein groups were combined in one category. Categories in orange of (**c**) are known to be regulated by or to regulate AMPK.

**Supplementary Figure 3.**
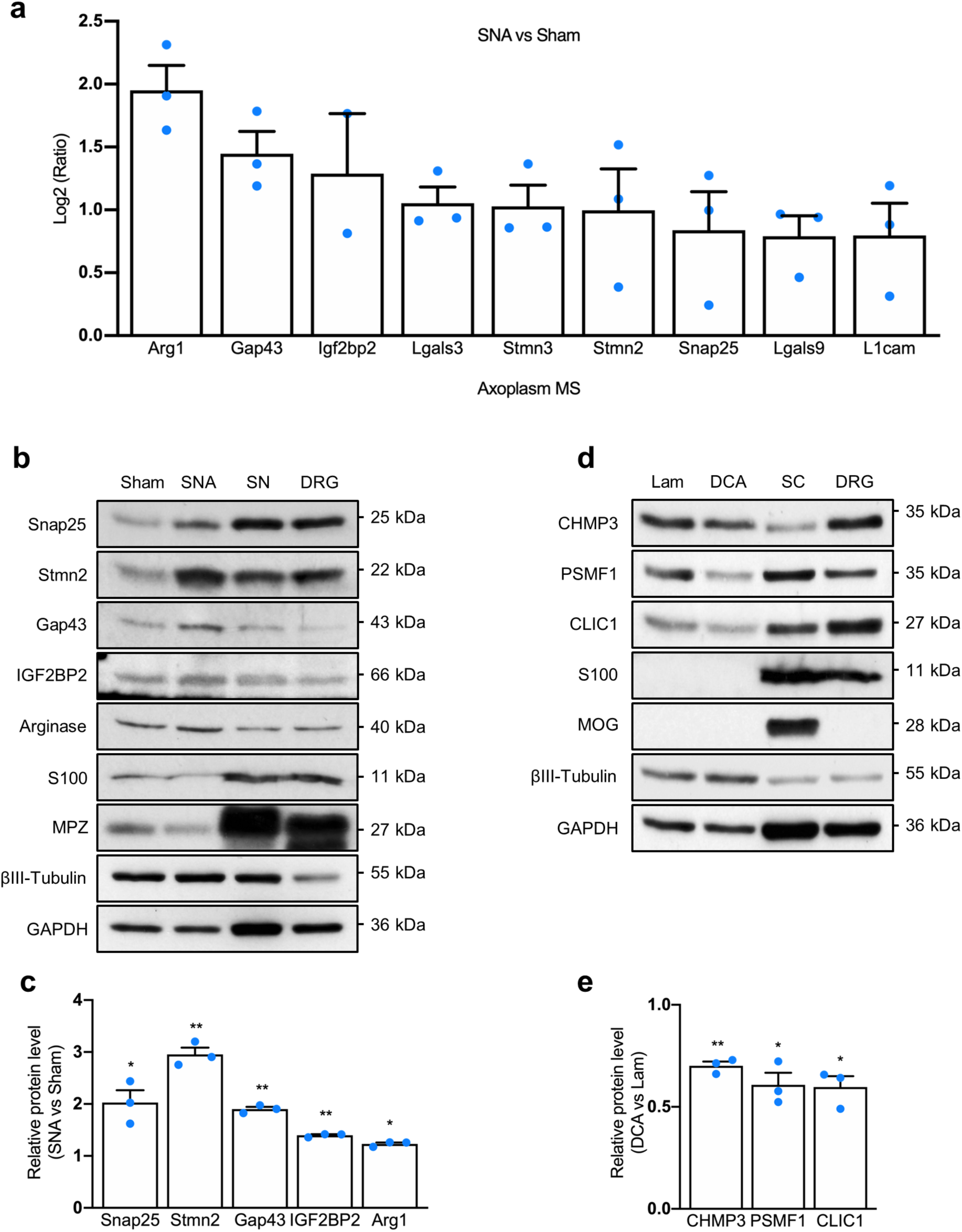
Regeneration associated genes and immunoblotting validation of axoplasmic protein expression identified by mass spectrometry. **a,** Bar graphs show axoplasmic proteins belonging to RAGs with FDR < 0.05, log2 ratio > 0.58 (SNA vs Sham) that are plotted in a log2 ratio scale. **b**, **c**, Immunoblots show validation of axoplasmic proteins identified by mass spectrometry after SNA vs Sham or DCA vs Lam (SN: sciatic nerve; SC: spinal cord). **d**, **e**, Bar graphs show quantification of immunoblots in (**b**) and (**c**) respectively. The expression level of each protein was quantified after normalization to GAPDH. Values represent means ± SEM (*p < 0.05; **p < 0.01; one-way ANOVA followed by Dunnett test).

**Supplementary Figure 4.**
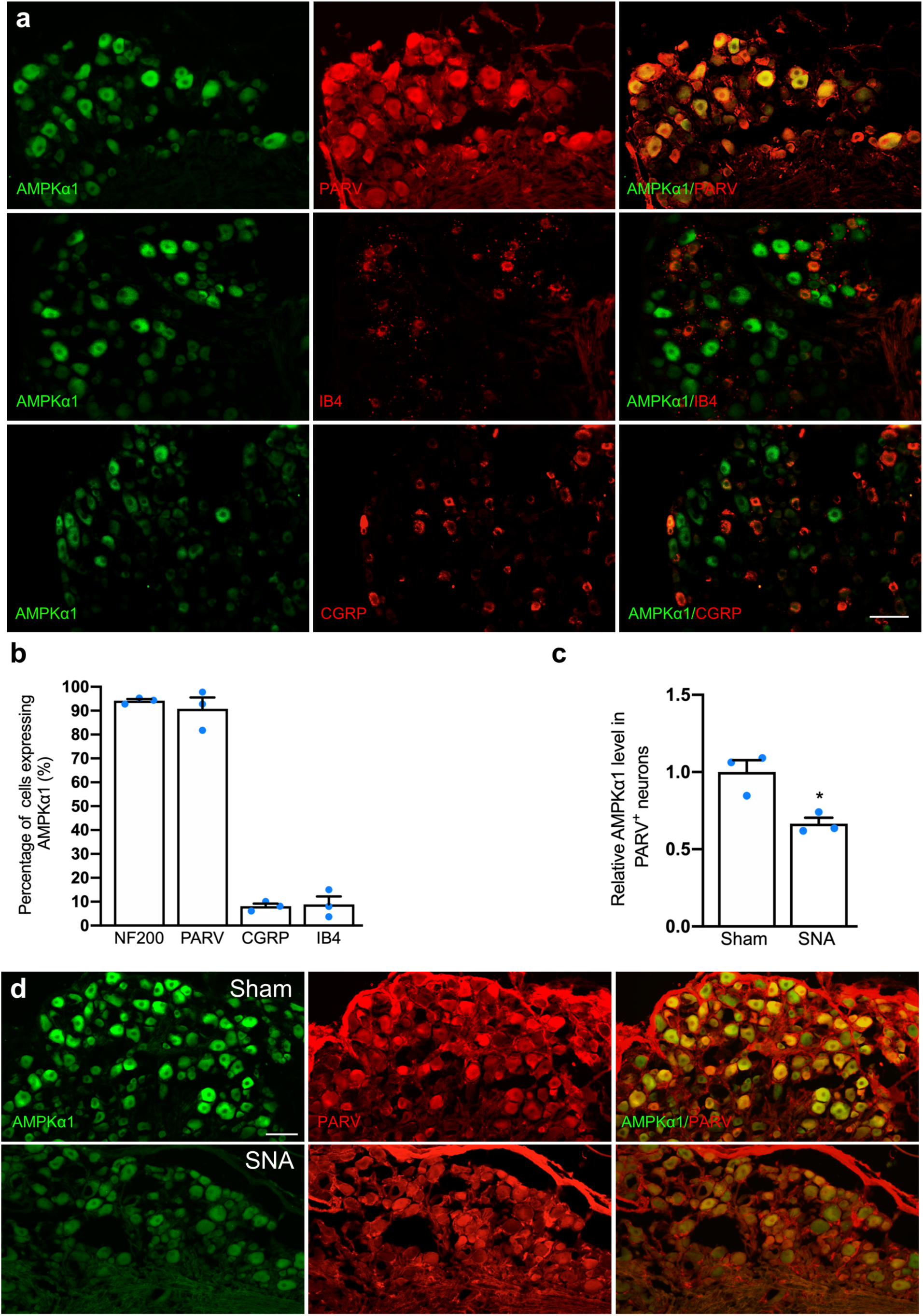
AMPKα1 expression in NF200, PARV, CGRP and IB4 positive DRG neurons. **a**, Representative fluorescence images of immunostaining for AMPKα1 and parvalbumin (PARV) or CGRP, or IB4^+^ in DRG neurons. Scale bar, 100 μm. **b**, Percentage of NF200, PARV, CGRP or IB4 positive neurons expressing AMPKα1. **c**, Quantification of immunostaining for AMPKα1 expression level of (**d**). n = 3 mice each group. The relative AMPKα1 expression level was quantified after normalization to the background (secondary antibody only). Values represent means ± SEM (*p < 0.05; two-tailed unpaired Student’s *t*-test). **d**, Representative fluorescence images of immunostaining for AMPKα1 and parvalbumin (PARV) in DRG neurons following Sham and SNA. Scale bar, 100 μm.

**Supplementary Figure 5.**
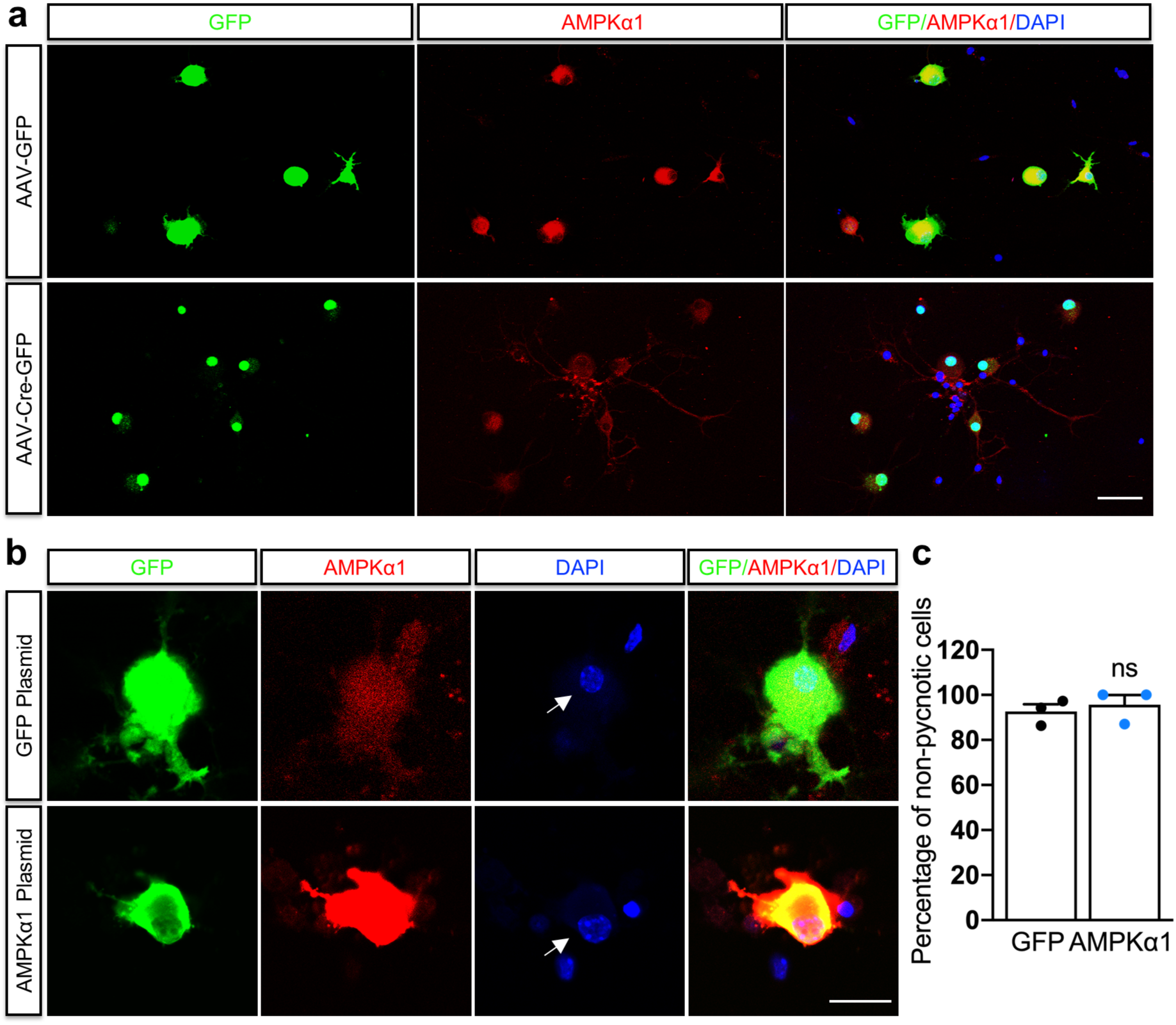
AMPKα1 expression in DRG neurons following conditional deletion or overexpression. **a**, Representative fluorescence images of immunostaining for GFP; AMPKα1 and DAPI in cultured DRG cells dissected from AMPKα1 floxed mice 48 h after transfection with AAV-GFP or AAV-Cre-GFP. Scale bar, 100 μm**. b**, Representative fluorescence images of immunostaining for GFP; AMPKα1 and DAPI in cultured DRG cells after electroporation with GFP plasmid or AMPKα1 plasmid at 48 h. Arrows show non-pycnotic DAPI positive nuclei. Scale bar, 50 μm**. c**, Quantification of the percentage of cells with non-pycnotic nuclei (**b**). Values represent means ± SEM (ns: not significant; two-tailed unpaired Student’s *t*-test).

**Supplementary Figure 6.**
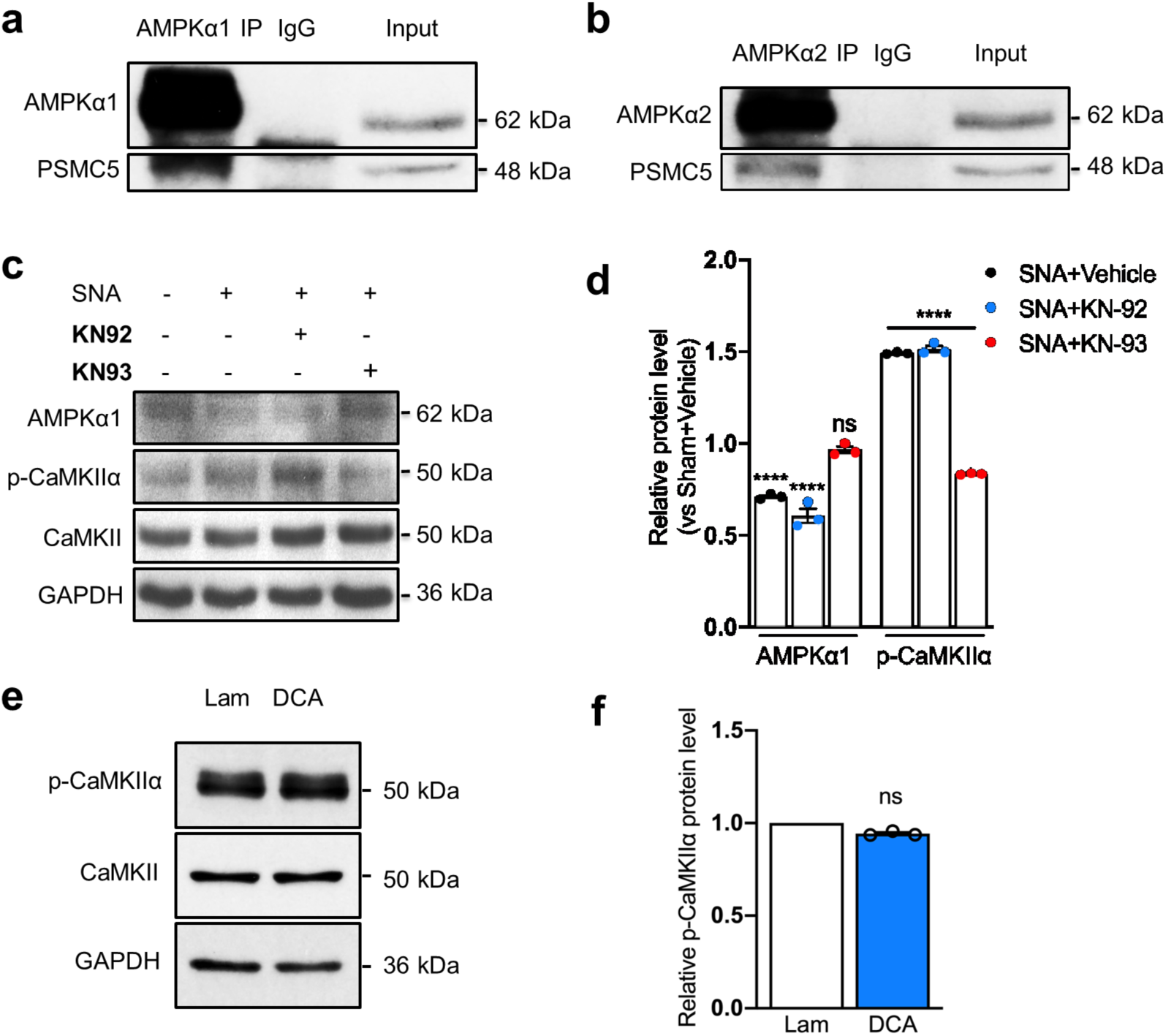
AMPKα1 and AMPKα2 interaction with PSMC5 and AMPKα1 expression after CaMKII inhibition. **a**, **b**, Immunoblots for PSMC5, AMPKα1 and AMPKα2 after AMPKα1 and AMPKα2 IP from DRG. **c**, Immunoblot shows AMPKα1 and p-CaMKIIα expression from sciatic DRG after intraperitoneal injection of the CaMKII inhibitor KN-93 (12.5 mg/kg) or KN-92 (an inactive derivative of KN-93). Sciatic DRGs were collected 24 h after SNA. **d**, Quantification of AMPKα1 and p-CaMKIIα expression level of (**c**). n = 3 independent experiments. Relative protein expression level (vs Sham+Vehicle) was quantified after normalization to GAPDH. Values represent means ± SEM (****p < 0.0001; ns: not significant; two-way ANOVA followed by Bonferroni test). **e**, Immunoblot shows p-CaMKIIα and CaMKII expression in sciatic DRG after Lam and DCA at 24 h. **f**, Quantification of p-CaMKIIα expression level of (**e**). n = 3 independent experiments. Relative protein expression level was quantified after normalization to GAPDH. Values represent means ± SEM (ns: not significant; two-tailed unpaired Student’s *t*-test).

**Supplementary Figure 7.**
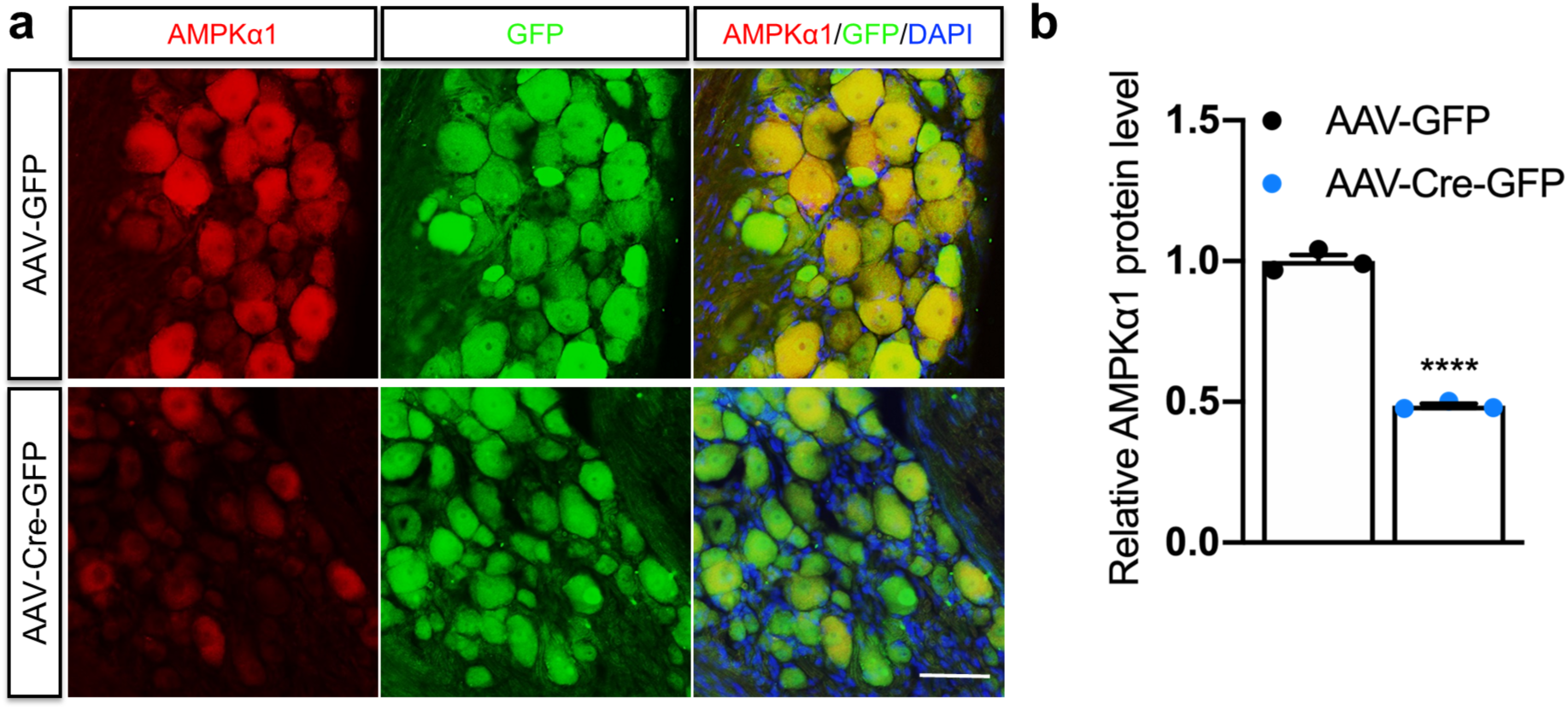
In vivo AMPKα1 conditional deletion in DRG neurons. **a**, Representative images of AMPKα1 and GFP immunostaining in DRG sections 5 weeks following AAV-GFP or AAV-Cre-GFP sciatic nerve injection. Scale bar, 50 μm. **b**, Quantification of AMPKα1 level of (**a**). n = 3 mice. Values represent means ± SEM (****p < 0.0001; two-tailed unpaired Student’s *t*-test).

**Supplementary Figure 8.**
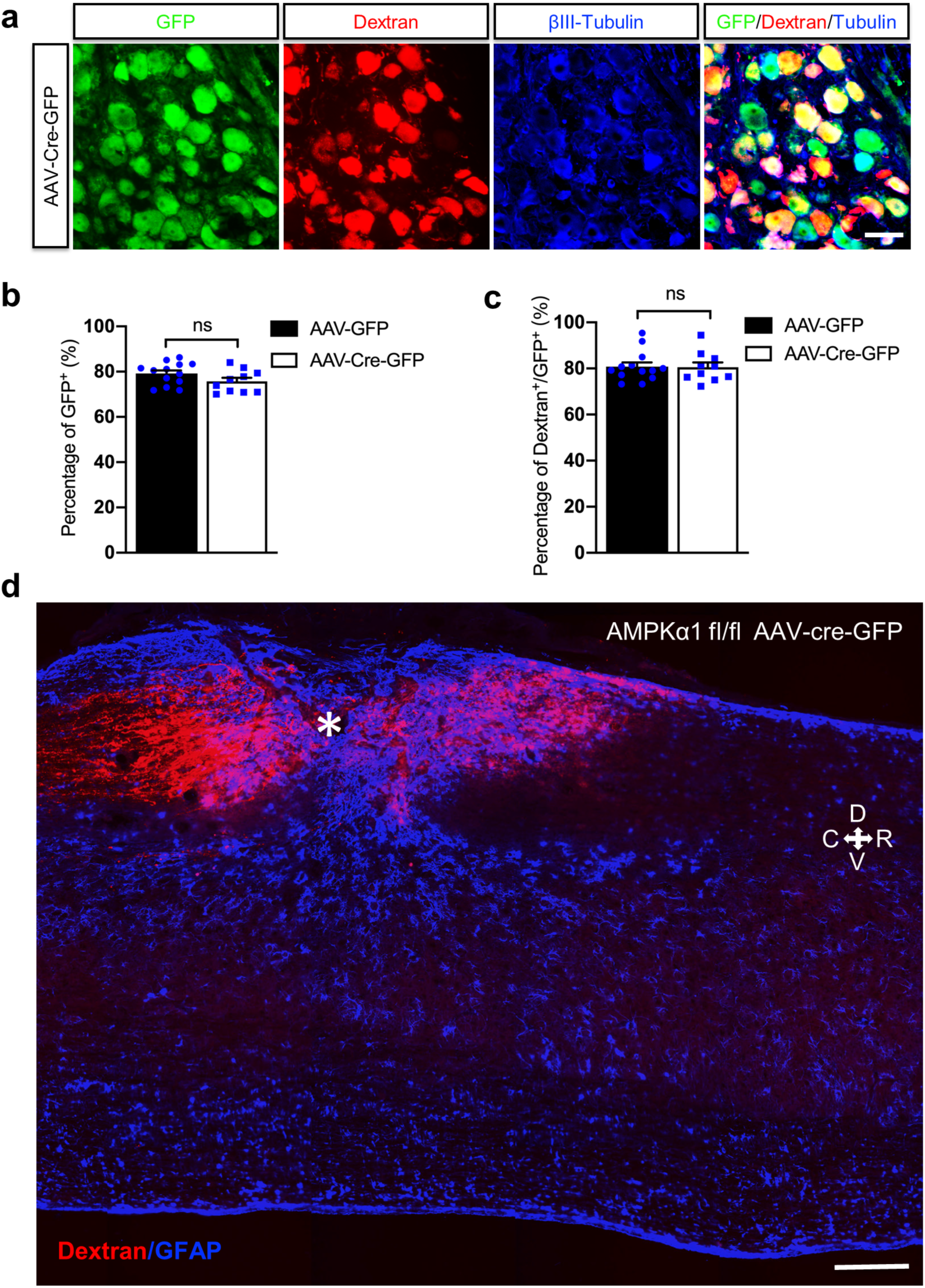
GFP and dextran co-localization in DRG neurons. **a**, Representative images of DRG sections from AAV-Cre-GFP sciatic nerve injected mice co-immunostained with antibodies against GFP and Dextran. Scale bar, 50 μm. **b**, **c**, Quantification of the percentage of GFP^+^ and Dextran^+^ / GFP^+^ cells following AAV-GFP or AAV-Cre-GFP. AAV-GFP, n = 13 mice; AAV-Cre-GFP, n = 10 mice. Percentage of GFP positive cells was calculated as the ratio of GFP^+^ versus TUJ1^+^ cells; percentage of Dextran^+^/GFP^+^ was calculated as the ratio of Dextran^+^/GFP^+^ versus TUJ1^+^ cells. Values represent means ± SEM (ns: not significant; two-tailed unpaired Student’s *t*-test). **d**, Longitudinal spinal cord section 5 weeks after SCI showing axonal labeling across the injured dorsal columns following deletion of AMPKα1. Dorsal column axons are labeled by sciatic nerve injected Dextran. Asterisk indicates the lesion epicenter. D; dorsal; V; ventral, C; caudal; R; rostral. Scale bar; 250 μm.

**Supplementary File 1.** Annotated list of proteins identified by axoplasmic proteomics in peripheral sciatic DRG axoplasm after sciatic peripheral injury-axotomy vs sham (SNA vs Sham) or in dorsal root axoplasm of sciatic DRG after central injury vs laminectomy (DCA vs Lam). Analyses of statistically significant proteins are included.

**Supplementary File 2.** Gene ontology (GO) and KEGG pathways of differentially enriched proteins in SNA vs Sham or DCA vs Lam sciatic DRG axoplasm.

**Supplementary File 3.** KEGG pathways of combined axoplasmic proteomics and DRG RNA sequencing (SNA vs Sham or DCA vs Lam).

**Supplementary File 4**. List of proteins identified by mass spectrometry following AMPKα IP including statistically differentially enriched proteins in SNA vs Sham.

